# Cognitive impairment and progressive neuroinflammation in mucopolysaccharidosis IIIA mice expressing the R245H Sgsh variant

**DOI:** 10.64898/2025.12.02.691757

**Authors:** Adeline A. Lau, Paul J. Trim, Helen Beard, Ashleigh Lake, Barbara M. King, Marten F. Snel, Kim M Hemsley

**Affiliations:** Childhood Dementia Research Group, College of Medicine and Public Health, Flinders Health and Medical Research Institute (FHMRI), Flinders University, Bedford Park, SA, Australia, 5042; Proteomics, Metabolomics and MS-Imaging Core Facility, South Australian Health and Medical Research Institute, Adelaide, Australia, 5000; Flinders Omics Facility, College of Medicine and Public Health, Flinders University, Bedford Park, SA 5042, Australia

**Author notes:** **Corresponding author:** Childhood Dementia Research Group, College of Medicine and Public Health, Flinders Health and Medical Research Institute (FHMRI), Flinders University, Bedford Park, SA, Australia, 5042.

## Abstract

Mucopolysaccharidosis (MPS) type IIIA (Sanfilippo syndrome) is an inherited childhood-onset dementia caused by insufficient SGSH enzymatic activity and subsequent accumulation of partially degraded heparan sulfate glycosaminoglycans. One of the most prevalent mutations is the Arg245His variant, representing up to 58% of mutations in some populations. Whilst other mouse models exist, none exhibit the mutations seen in humans, thus their amenability to testing of therapeutics such as pharmacological chaperones, which are often mutation-dependent, is limited. We have used CRISPR/Cas 9 gene editing to generate the first *Sgsh*^R245H^ knock-in mouse model of MPS IIIA, which subsequently underwent extensive phenotyping. *Sgsh*^R245H^ mouse brain exhibited progressive accumulation of heparan sulfate, endo/lysosomal system expansion and elevated levels of GFAP-reactive astroglial staining and activated microglia. Significant perturbation in several novel lipid species was observed in the CA1 region of the hippocampus using Matrix-Assisted Laser Desorption/Ionisation Mass Spectrometry Imaging. Spatial memory deficits were apparent in the Morris Water Maze probe and Y-maze tests, and Elevated Plus Maze exploration revealed reduced anxiety-like behaviours. *Sgsh*^R245H^ MPS IIIA mice recapitulate key characteristics of the human disorder and represent a useful tool for studying disease pathogenesis and evaluating novel therapeutic approaches.

**Graphical Abstract:** 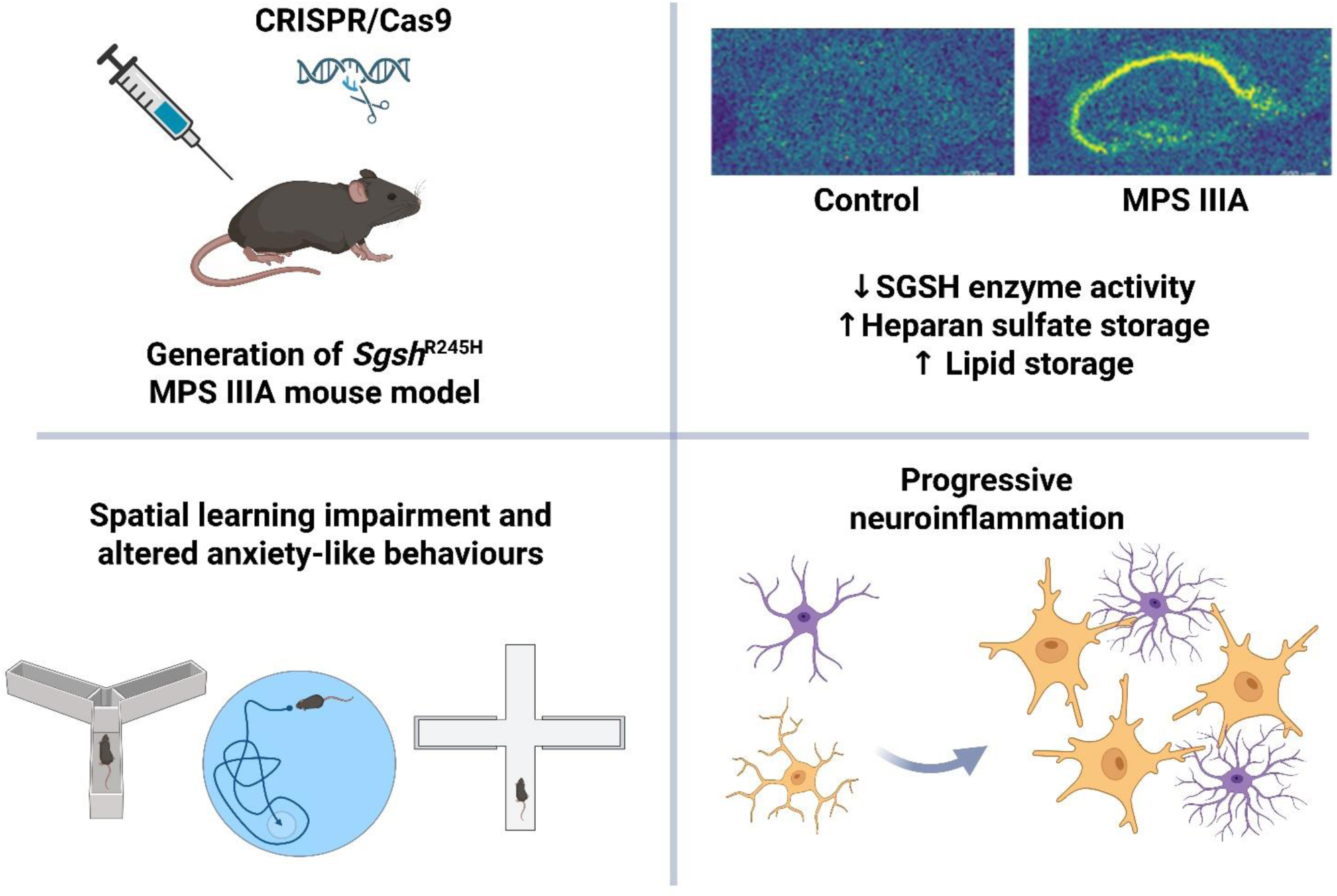

## Introduction

Mucopolysaccharidosis (MPS) types IIIA-D are rare, neurodegenerative lysosomal storage disorders that arise from insufficient activity of one of four specific enzymes involved in heparan sulfate (HS) catabolism. In the case of MPS IIIA (OMIM #252900), a lack of *N*-sulfoglucosamine sulfohydrolase (SGSH; EC 3.10.1.1) causes clinical signs that frequently begin in toddlerhood and are predominantly neurological, e.g., hyperactivity; the progressive loss of learned skills, vision, hearing, and mobility; and sleep disturbance (1). The median age of death is 18 years (2), and there are no approved therapies. However, several approaches have been investigated in clinical trials including the delivery of replacement recombinant SGSH enzyme either in its unmodified form or via chemical modification to allow the enzyme to cross the blood-brain barrier (clinicaltrials.gov identifier NCT01155778; NCT03811028), gene transfer of adeno-associated virus serotypes-9 or rh10 vectors, or *ex vivo* lentiviral-mediated gene therapy via haematopoietic stem cell transplantation (NCT01474343; NCT02716246; NCT04201405).

Whilst we and others have utilised the spontaneous D31N *Sgsh* mouse model of MPS IIIA (3, 4), or created new models of MPS IIIA, e.g., conditional *Sgsh* knockout (5, 6) or compound mutants (7–9) to study disease pathogenesis and evaluate therapeutic approaches, MPS IIIA mice carrying common human missense mutations have not been generated to date. This is despite missense mutations occurring in the vast majority of MPS IIIA patients (∼76%; 10, 11, 12). The R245H *SGSH* mutation, where an arginine residue in the helix alpha-7 loop is converted to a histidine, is a high-frequency allele representing up to 58% (The Netherlands), 41% (Australia), and 35% (Germany) of the MPS IIIA patient population (10, 13). It is also prevalent in the Cayman Islands, potentially due to a founder effect, with 47% of the 77 tested individuals from one family identified as carriers (14). The R245H *SGSH* mutation reduces the amount of normal SGSH protein produced to 0.05% (15) and is associated with the classic early-onset, rapidly-progressing phenotype (2).

One therapeutic approach under investigation for lysosomal storage disorders is pharmacological chaperone therapy. Pharmacological chaperones are often mutation-specific in their action, necessitating testing of compounds against individual patient mutations (16). The available MPS IIIA mouse models are unsuitable for testing clinically relevant pharmacological chaperones. Further, recent studies in MPS IIIC mice have indicated a toxic gain of function in missense mutants due to misfolded HGSNAT (17). Whether misfolded human SGSH has a similar impact is unknown. To begin to address these two knowledge gaps, we report the generation and characterisation of transgenic mice expressing biallelic R245H *Sgsh* mutations.

## Materials and Methods

### Animal ethics and biosafety approvals

Animal ethics approval was granted by Flinders University in accordance with the 2013 National Health and Medical Research Council’s Australian Code For The Care And Use Of Animals for Scientific Purposes, 8^th^ edition (approval number 5726). Mice were housed in an enriched environment with a 12h:12h light/dark cycle and had *ad libitum* access to chow and water. Institutional biosafety approval was also granted (#2022-21).

### Generation of Sgsh^R245H^ mutant mice

CRISPR/Cas9 technology was used by the SA Genome Editing Facility (Adelaide, Australia) to generate *Sgsh*^R245H^ mutant mice. The CRISPR guide sequence 5’ CTCAGTACACCACCATCGGG was used. A prime editing reverse transcribed guide sequence was used as a donor template to convert the CGG arginine to a CAT histidine at codon position 245 (**Fig. 1a**). C57BL/6 mouse zygotes were microinjected with 50ng/µL Sgsh prime editing gRNA and 150ng/µl PEA1 mRNA (generated from modified plasmid Addgene #171991) and implanted into pseudopregnant females. Genomic DNA was extracted from mouse pups using Roche High pure PCR template preparation kits, for screening. An amplicon of 140 bp containing the putative mutation was amplified by PCR (forward primer 5’TCATCCTCAGGTGCCCTACT and reverse primer 5’ GGTCCCTGGCCTATGATGAG) and digested with *Nde*I restriction enzyme. Mutant *Sgsh*^R245H^ mice revealed digestion products of 81 and 59 bp, whereas the wild-type mice were undigested (140 bp). Two founder males were each bred with three wild-type C57BL/6 females to establish separate pedigreed breeding colonies. Sanger sequencing confirmed the genotypes of the founders and F1 mice. *Sgsh*^+/+^, *Sgsh*^+/R245H^ and *Sgsh* ^R245H /R245H^ are henceforth referred to as wild-type, heterozygote and MPS IIIA. The *Sgsh*^R245H^ MPS IIIA mouse line has been provided to the Australian Phenome Bank (strain #10512) for cryopreservation and sharing with the scientific community.

**Figure 1:**
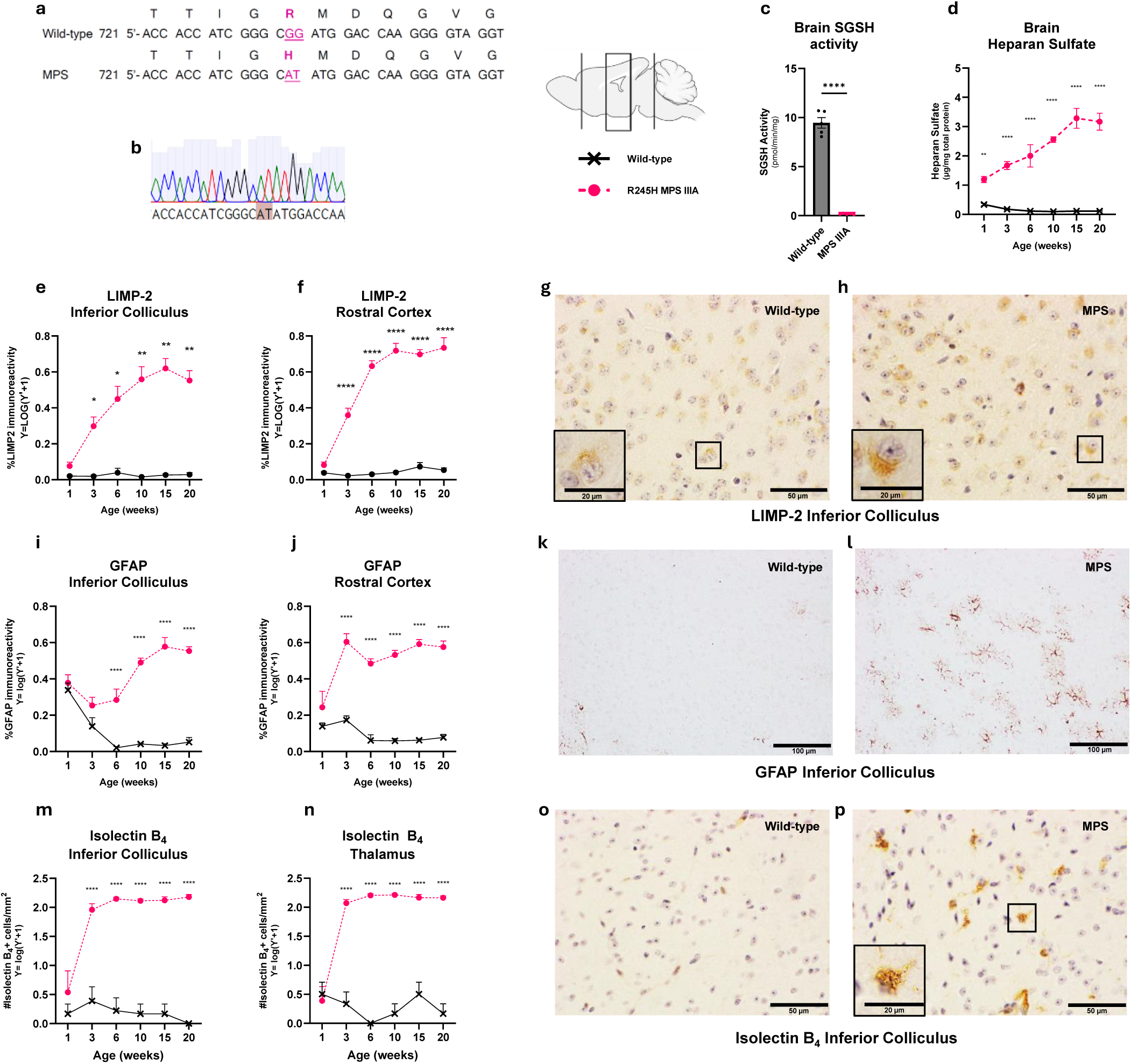
Generation of the *Sgsh*^R245H^ variant in mice by germline CRISPR/CasG gene targeting, with subsequent evaluation of biochemical and pathological features. (a) Nucleotide and amino acid sequence alignments of the murine *Sgsh* wild-type and mutants highlighting the conversion of an arginine to a histidine residue in R245H *Sgsh* mutant mice. (b) Sanger sequencing results confirmed the nucleotide sequence change in transgenic MPS IIIA mice. (c) Fluorometric SGSH assays on brain homogenates, as indicated by the rectangle in the schematic sagittal section, confirmed the reduction in SGSH enzymatic activity. (d) Total HS disaccharide content was quantitated via tandem mass spectrometry. SGSH activity and HS content were normalised to total protein content. (e-p) Immersion- fixed mouse brain tissue from wild-type or MPS IIIA mice, was sectioned in the sagittal plane and stained using immunohistochemical/histochemical methods. (e-h) Lysosomal integral membrane protein-2 (LIMP2) staining as a marker of endosomal/lysosomal compartment size was measured in the (e,g,h) inferior colliculus and (f) rostral cerebral cortex. (i-l) The percentage of thresholded GFAP reactivity showed elevated levels of astrogliosis in the (I,k,l) inferior colliculus or (j) cerebral rostral cortex of MPS IIIA mice. (m-p) Increased numbers of isolectin B_4_-positive microglia per tissue area were quantified in the (m,o,p) inferior colliculus and (n) thalamus of MPS IIIA mice. Representative images of (g,h) LIMP-2, (k,l) GFAP or (o,p) isolectin B_4_ staining from 20-week-old wild-type and MPS IIIA mice in the inferior colliculus are shown. Scale bars are (g,h,o,p) 50µm, inset 20µm, or (k,l) 100µm. n=5 mice/group; mixed sexes. \**p*≤0.05; \*\**p*≤0.01; \*\*\*\**p*≤0.0001.

### Genotyping

Genomic DNA was extracted from mouse ear tags by heating at 100°C in 10% Chelex resin for 20 minutes. After centrifugation at 13,000 rpm, the DNA was quantitated on a Nanodrop One and diluted in water to a final concentration of 2.5 ng/µl. Mice were genotyped using custom Taqman® SNP genotyping assays that contained primers flanking the mutation (forward primer 5’-CAGACCTAGCTGCTCAGTACAC and reverse primer 5’- AGGGACAGCCTCAGTCACT) and allele-specific probes with non-fluorescent quenchers at the 3’ end (wild-type probe 5’-CTTGGTCCATCCGCCCGA-VIC dye; Sgsh-R245H probe 5’- CCTTGGTCCATATGCCCGA-FAM dye) (Thermofisher Scientific; 4332077).

Reactions were performed in 384-well PCR plates with each well containing 1xTaqMan® Genotyping Master Mix (Thermofisher Scientific; 4371355), 1x of the custom SNP assay and 2.5 ng of DNA (5 μL total volume per well). The plates were sealed and amplification performed on a Biorad CFX Opus 384 Real-Time PCR System using the following cycling conditions: 60°C for 30s pre-read stage; 95°C 10 min hold stage; 40 cycles of 95°C for 15s and then 60°C for 1 min PCR amplification stage; 60°C for 30s post-read stage. Data was analysed using Biorad CFX Maestro Software (v2.0).

#### Tissue collection

For *ex vivo* assays, one-week-old mice were euthanized by decapitation. Mice that were ≥3 weeks old were humanely killed by slow-fill carbon dioxide inhalation. Following intra-cardiac perfusion with PBS, the brain was removed, dissected along the longitudinal fissure and the left hemisphere post-fixed in 4% paraformaldehyde in PBS for 2 days. The right brain hemisphere was further dissected into five 2-mm hemi-coronal slices and snap frozen in liquid nitrogen, with slices 1 and 5 containing the olfactory bulb and cerebellum, respectively. For MALDI mass spectrometry imaging, 20-week-old mouse brain was snap frozen in hydrogel (7.5% HMPC with 2.5% PVP (18)) using dry ice-cooled isopentane. Brain tissue was cryo-sectioned at 10 μm in the sagittal plane onto indium-tin-oxide (ITO)-coated conductive microscope slides (Bruker) and stored at −80°C until imaging.

### Heparan sulfate quantitation

Brain slice 3 was transferred to Lysing Matrix D tubes (MP Biomedicals; 116913500) containing 20 mM Tris, 0.5 M NaCl, pH 7.4. Tissues were homogenised at 4000 rpm for 10 seconds on a BeadBug Benchtop Homogeniser (Pathtech). The total protein content was determined by MicroBCA (Thermofisher Scientific; 23235) and 10 µg total protein lyophilised. HS was measured according to the method of He *et al*., (19) except butanolysis used 25 μL of 2,2-dimethoxypropane and 500 μL of 3 M HCl in butan-1-ol. Liquid chromatography tandem mass spectrometry was performed on an API4000 QTrap (ABSciex, Concord, Ontario, Canada) coupled to a Waters Acquity UPLC system. Peak area ratios were determined using Analyst 1.6.2 software (ABSciex).

### SGSH activity

Brain homogenates prepared as above, were dialysed into 200 mM sodium acetate, pH 6.5 overnight at 4°C. SGSH enzymatic activity was assayed as described previously (20) using 10 µL homogenate and 20 µL 5 mM substrate (4-methylumbelliferyl 2-deoxy-2-sulfamino-a-D-glucopyranoside sodium salt; Carbosynth EM06602) at 47°C for 16 hours. A 4-methylumbelliferone (Sigma; M1381) standard curve was included to convert sample counts to SGSH activity rates in units of pmol/min. Fluorometric counts were determined on a Molecular Devices Spectramax iD5 microplate reader at excitation λ355 nm and emission λ460 nm.

### Histology

The left-brain hemisphere was processed into paraffin. Six-micron-thick sections were cut on a rotary microtome (HM325, Thermofisher Scientific) at ∼0.6-0.72-mm lateral to the midline (21), and mounted on Superfrost^TM^ Plus glass slides (Thermofisher Scientific). Immunohistochemical and histochemical staining were performed according to previously published methods (22). Heat-mediated epitope retrieval in citrate buffer, pH6.0, [lysosomal integral membrane protein-2 (LIMP2) and glial fibrillary acidic protein (GFAP)] or 1 mM EDTA, pH8.0 (peroxidase-conjugated isolectin B_4_) was performed before overnight antibody/lectin incubation (reagent details in **Supp. Table 1**). Species-specific biotinylated secondary antibodies and Vectastain Elite ABC kit reagents (PK-6100; Vector Laboratories) were applied prior to chromogenic visualisation using diaminobenzidine metal-enhanced substrate (#11718096001; Roche).

Staining was batched and analysis was carried out by a researcher blind to age/genotype. Sections were viewed on an Olympus BH2 microscope fitted with an Olympus DP22 camera. Images were captured using Olympus CellSens imaging software. LIMP2 and GFAP analysis was based on the optical density of positive staining, using FIJI software (23) and reported as a percent positive immunoreactivity. The number of isolectin B_4_-stained cells was counted manually per unit area of brain and reported as the number of activated microglia/mm^2^.

#### Matrix-Assisted Laser Desorption/Ionisation Mass Spectrometry Imaging (MALDI-MSI)

Slides were thawed in a vacuum desiccator for 60 minutes prior to opening to ensure no condensation. Samples were prepared for negative ion imaging using a modified method from Yang *et al*., (24). Briefly, slides were washed (30 sec) in 50 mM ammonium formate solution, dried and spray-coated with Norharmane matrix using a SunCollect MALDI sprayer (SunChrom GMBH.) Matrix was applied in 15 layers of 7.5 mg/mL norharmane in MeOH. Flow rates were 10, 20, 30 and 40 µL/minute for layers 1, 2, 3 and 4-15, respectively. Samples were stored at −20°C for 24 hours before imaging. MALDI-MSI was conducted using a timsTOF fleX mass spectrometer (Bruker), the instrument was mass calibrated prior to analysis using sodium formate calibration solution. Negative ion mode imaging was performed using a 20 µm pixel size, over a range of 300-3000*m/z*. Data were analysed using SCiLS Lab MVS Pro (2026a). Images were normalised to the Root Mean Square. Lipid analysis was performed by searching an in-house curated lipid database containing 390 commonly observed negative lipid ions (**Supp. Table 2**). A more expansive ganglioside database of theoretical masses was produced by downloading the LipidMaps LMSD-ganglioside database using [M-H]^-^ ions with sum composition. Peak areas were calculated within +/- 15 ppm of the theoretical *m/z* for an ROI within the CA1a/b region of the hippocampus (21). Lipid masses with a mean intensity < 40 a.u. were removed before statistical analysis. Statistically different (*p*<0.05) lipid masses were manually curated to ensure accurate mass and removal of isotopic interfering peaks.

#### Behavioural test battery

The mice were separated into single housing at 22-weeks of age and acclimatised to the behaviour testing room for >4 days before the first test. Experimental mice were divided into two cohorts with equal numbers of genotypes/sexes, and then randomly assigned a code and order within each group to allow blinded testing (n=16/group). Behavioural tests were conducted in a pre-determined order by one experimenter (AL) using published methods. Additional details of the behavioural phenotyping are described in the **Supplementary Information**. In brief, mice were tested in the Morris Water Maze [cohort 1, 22-weeks-old; cohort 2, 23-weeks-old; (25)]); open field locomotor test [24-weeks-old (26)]; Elevated plus maze [24-weeks-old (26)]; Y-maze test [25-weeks-old (27)] followed by a motor test battery [25-weeks-old (5, 28)] consisting of neuromuscular grip strength, pole test, negative geotaxis, and gait. Videos were recorded for some tests using a Logitech BRIO camera and Logitech capture software for offline analysis using ANYMaze software. Except for the Morris water maze and gait corridor, each apparatus was cleaned with 70% ethanol between trials.

### Statistics

Data are presented as the mean+SEM. Statistical analyses were undertaken using GraphPad Prism 10, SPSS for Windows 28.0.1.1(15) or Metaboanalyst 6.0. Repeated measures two-way ANOVA was used to determine statistical significance with genotype and time as the independent factors (growth curves or Morris Water Maze acquisition phase). Two-way ANOVA (genotype x age; genotype x colony) or unpaired t-tests were used for other assays. The Bonferroni *post-hoc* test was applied to adjust for multiple comparisons between groups. *p*≤0.05 was regarded as statistically significant. MALDI-MSI data were log-transformed (base 2) and analysed using principal component analysis and two-way ANOVA with FDR *post-hoc* correction.

## Results

### Generation of Sgsh^R245H^ mutant mice via CRISPR/Cas3 editing technology

CRISPR/Cas9 gene editing technology introduced a histidine in place of the arginine residue corresponding to the R245H mutation in the *Sgsh* gene of C57BL/6 mice (**Fig. 1a**). Two founder males were each mated with wild-type C57BL/6 females to establish independent, pedigreed, congenic breeding colonies. Sanger sequencing confirmed the genotypes of the founder mice and F1 generation heterozygote progeny (**Fig. 1b**). We developed a custom TaqMan single-nucleotide polymorphism (SNP) genotyping method to improve the assay throughput. Genomic DNA was extracted from an ear clip and amplified in the presence of a primer pair flanking the mutation and VIC- or FAM-dye labelled DNA probes that specifically hybridise to either the wild-type or R245H mutant *Sgsh* allele sequence. Allelic discrimination of wild-type, heterozygote, or R245H *Sgsh* status could be determined in <2.5 hours (**Supp. Fig. 1**).

One colony was selected for expansion (founder male SPAT/5.2.24). Mutant female and male mice were fertile, and various genotype pairing combinations had similar litter sizes and pair-to-birth intervals (**Supp. Table 3**). Progeny from heterozygous inter-cross pairs demonstrated the expected Mendelian frequencies (27.8% wild-type, 49.6% heterozygote, and 22.5% MPS IIIA). Weekly body weight measurements showed that both female and male MPS IIIA mice were significantly heavier than their sex- and age-matched unaffected counterparts from 6 weeks of age (**Supp. Fig. 1**).

### Low SGSH activity and progressive HS accumulation in R245H MPS IIIA mice

SGSH activity was measured using 4-methylumbelliferyl 2-deoxy-2-sulfamino-α-D-glucopyranoside substrate (20). R245H MPS IIIA mouse brain homogenates had greatly reduced SGSH activity, with ∼1.7% of that measured in wild-type mice at 20 weeks of age (**Fig. 1c**). We then quantified HS levels in affected mice using a liquid chromatography tandem mass spectrometry method (19). Brain tissue was harvested from wild-type and MPS IIIA mice aged between 1- and 20-weeks of age. Total HS disaccharide concentrations were significantly elevated in MPS IIIA mouse brain at all ages assessed compared to age-matched wild-type controls (genotype *p*<0.0001; **Fig. 1d**). Progressive accumulation of HS was evident in MPS IIIA mouse brain, increasing from 3.5-fold normal levels at 1-week of age to 27-fold normal at 20- weeks (age *p*<0.0001).

Expansion of the endo/lysosomal system in brain was examined using immunohistochemical staining of LIMP2, a component of late endosomal and lysosomal membranes. LIMP2 staining was visualised as a punctate cytoplasmic signal and representative images of the staining pattern are presented in **Fig. 1g,h**. Low levels of LIMP2 staining were measured in the inferior colliculus and rostral cerebral cortex of wild-type mice at all ages examined (**Fig.1e,f**). One-week-old unaffected and MPS IIIA mice had similar LIMP2 levels. However, a progressive increase in LIMP2 immunoreactivity was evident in MPS IIIA mice (age and genotype *p*<0.0001 for both brain regions; **Fig. 1e,f**).

### Neuroinffammatory markers increase with age in R245H *Sgsh* MPS IIIA mice

GFAP levels were measured in MPS IIIA and age-matched control brain sections as a marker of astrocytic activation (**Fig. 1i-l**). Low levels of GFAP-positive immunohistochemical staining were observed in wild-type mouse brains at all ages examined. There was no difference in GFAP staining between wild-type and affected mice at 1 week of age. From 3 and 6 weeks of age, GFAP reactivity was significantly higher in MPS IIIA mice in the rostral cortex and the inferior colliculus, respectively. We also measured microglial activation using isolectin-B_4_-reactivity (**Fig. 1m-p**). Compared to the number of activated microglia counted in wild-type mouse brain, MPS IIIA mice showed a significantly elevated number of activated microglia in the thalamus and inferior colliculus from 3 weeks of age. This increased number of microgliosis was sustained until 20 weeks of age, the last time-point of the study.

### MALDI-MSI screening revealed secondary lipid elevations in the MPS IIIA CA1 hippocampus

To interrogate the brain lipid landscape of *Sgsh*^R245H^ MPS IIIA mice, we employed a MALDI-MSI approach, utilising the well-characterised spontaneous *Sgsh*^D31N^ MPS IIIA mouse model (4) as a comparative benchmark. The CA1 hippocampus was selected as a region of interest, owing to its functional roles in motor learning and memory processing (29). PERMANOVA of principal component analysis (PCA) scores indicated significant genotype-associated separation in PC3-PC5 space (R^2^=0.35, p=0.039), despite weaker separation in higher-variance components (PC1-PC2, R^2^=0.15, p=0.20; PC1-PC3, R^2^=0.24, p=0.090; PC3-PC4, R^2^=0.24, p=0.070), indicating genotype effects are robust, but captured in lower-variance components (**Fig. 2a**). Subsequent univariate ANOVA revealed a total of 20 statistically significant genotype-associated *m/z* features (p<0.05). Statistically significant lipid masses were each assessed manually and removed if the peak selected for integration was not centred within the theoretical mass window, and were also removed if there was isobaric interference from an isotope of another mass feature contributing >50% of the observed intensity, generating 11 high-confidence *m/z* features, with no colony or interaction differences observed (**Fig. 2a**; **Supp. Table 4**). Tentative lipid identifications based on accurate mass included: *m/z* 806.5464 (SHexCer 36:1;O2); *m/z* 820.5256 (SHexCer 36:2;O3); *m/z* 822.5413 (SHexCer 36:1;O3); *m/z* 865.5031 (bis(monoacylglycero)phosphate [BMP] 44:12); *m/z* 867.5188 (BMP 44:11); *m/z* 968.5992 (SHex2Cer 36:1;O2); *m/z* 1179.7372 (monosialodihexosylganglioside [G_M3_] 36:1); *m/z* 1207.7685 (G_M3_ 38:1); *m/z* 1382.8166 (monosialotrihexosylganglioside [G_M2_] 36:1); *m/z* 1410.8479 (G_M2_ 38:1); and *m/z* 1470.8326 (disialodihexosylganglioside [G_D3_] 36:1) (**Table 1**; **Fig. 2b,c; Supp. Figs. 2-13**). No significant hits were observed in positive ion mode (data not shown).

**Figure 2:**
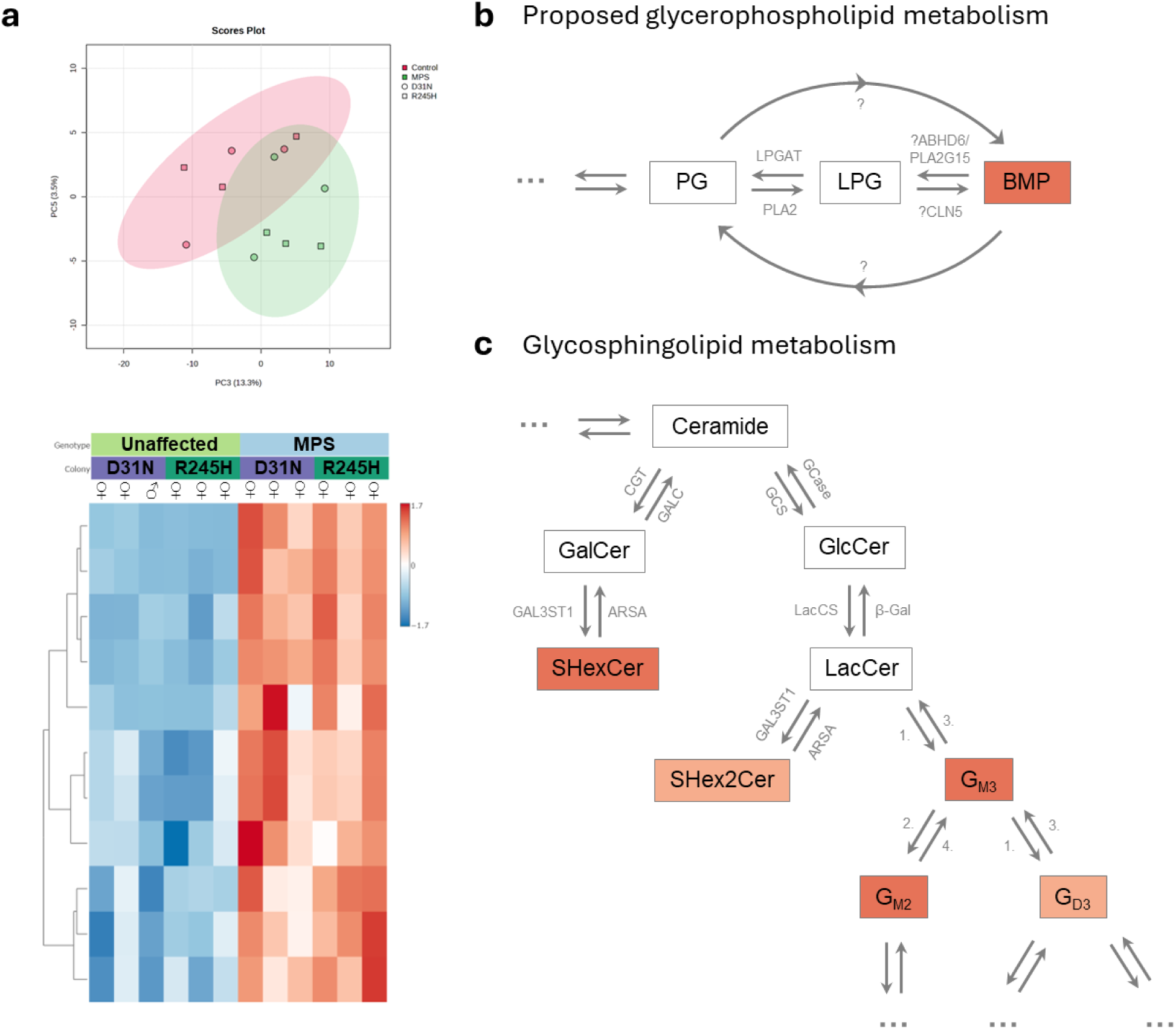
Identification of elevated *m/z* lipid features in 20-week-old MPS IIIA mouse hippocampus with MALDI-MSI. (a) Principal component analysis scores plot depicting significant genotype variation in the PC3-PC5 space and corresponding univariate ANOVA heatmap of 11 significant *m/z* features in MPS IIIA (n=3 *Sgsh*^D31N^ C n=3 *Sgsh*^R245H^) mice compared to unaffected controls (n=3 *Sgsh*^D31N^ C n=3 *Sgsh*^R245H^) at 20 weeks. (b) Proposed metabolic pathway of bis(monoacylglycero)phosphate (BMP) via phosphatidylglycerol (PG) and lysophosphatidylglycerol (LPG) glycerophospholipids. (c) Simplified depiction of glycosphingolipid metabolism, beginning with the hexose addition to ceramide to form either galactosylceramide (GalCer) or glucosylceramide (GlcCer). GalCer is further metabolised to SHexCer sulfatide, whereas GlcCer is converted to lactosylceramide (LacCer) and subsequently metabolised to SHex2Cer sulfatide or gangliosides. Highlighted glycerophospholipids and glycosphingolipids show increased levels in MPS IIIA mice, with darker shading reflecting elevation across multiple sub-species within each lipid class. G_M3_, monosialodihexosylganglioside; G_M2_, monosialotrihexosylganglioside; G_D3_, disialodihexosylganglioside; LPGAT, lysophosphatidylglycerol acyltransferase; PLA2, phospholipase A2; ABHD6, alpha/beta hydrolase domain-containing 6; PLA2G15, phospholipase A2 group 15; CLN5, ceroid lipofuscinosis neuronal protein 5; CGT, galactosylceramide synthase; GALC, galactosylceramidase; GCS, glucosylceramide synthase; GCase, beta glucocerebrosidase; GALST1, galactose-3-O- sulfotransferase 1; ARSA, arylsulfatase A; LacCS, lactosylceramide synthase; β-Gal, β-galactosidase; ?, unelucidated/theoretical mechanism/enzyme; …, further metabolism. Enzymatic steps: (1) sialyltransferase, (2) galactoside sialytransferase, (3) neuraminidase, (4) hexosaminidase.

**Table 1:** Mass-to-charge (*m/z*) features significantly altered in MPS IIIA mice and their putative identities. Mean signal intensities (n = 3 biological replicates) were normalised to root mean square. Fold change represents the ratio of mean signal intensity in MPS IIIA mice relative to unaffected controls for both *Sgsh*^R245H^ and *Sgsh*^D31N^ colonies. Significance is reflected as *p≤0.05, **p≤0.01, ***p≤0.0001, and ****p≤0.00001. A.U., arbitrary units; BMP, bis(monoacylglycero)phosphate; G_M3_, monosialodihexosylganglioside; G_M2_, monosialotrihexosylganglioside; G_D3_, disialodihexosylganglioside; MPS, mucopolysaccharidosis IIIA.

| *m/z* | Putative Identity | Mean Signal Intensity/Root Mean Square (A.U.) |  |  |  | Fold Change |  | Genotype Significance |
| --- | --- | --- | --- | --- | --- | --- | --- | --- |
|  |  | Unaffected |  | MPS |  | *Sgsh*<sup>R245H</sup> | *Sgsh*<sup>D31N</sup> |  |
|  |  | *Sgsh*<sup>R245H</sup> | *Sgsh*<sup>D31N</sup> | *Sgsh*<sup>R245H</sup> | *Sgsh*<sup>D31N</sup> |  |  |  |
| 806.5464 | SHexCer 36:1;O2 | 293.8 | 312.3 | 511.9 | 601.3 | 1.7 | 1.9 | * |
| 820.5256 | SHexCer 36:2;O3 | 47.6 | 50.5 | 67.6 | 67.8 | 1.4 | 1.3 | * |
| 822.5413 | SHexCer 36:1;O3 | 224.8 | 240.3 | 459.0 | 531.2 | 2.0 | 2.2 | * |
| 865.5031 | BMP 44:12 | 120.3 | 109.3 | 203.4 | 176.0 | 1.7 | 1.6 | * |
| 867.5188 | BMP 44:11 | 47.4 | 43.1 | 63.0 | 59.1 | 1.3 | 1.4 | * |
| 968.5992 | SHex2Cer 36:1;O2 | 40.4 | 44.6 | 139.9 | 143.1 | 3.5 | 3.2 | ** |
| 1179.7372 | G<sub>M3</sub> 36:1 | 58.8 | 55.5 | 180.2 | 165.3 | 3.1 | 3.0 | ** |
| 1207.7685 | G<sub>M3</sub> 38:1 | 13.8 | 13.6 | 20.8 | 20.9 | 1.5 | 1.5 | * |
| 1382.8166 | G<sub>M2</sub> 36:1 | 30.9 | 31.9 | 228.0 | 231.8 | 7.4 | 7.3 | **** |
| 1410.8479 | G<sub>M2</sub> 38:1 | 13.6 | 13.7 | 22.8 | 27.9 | 1.7 | 2.0 | * |
| 1470.8326 | G<sub>D3</sub> 36:1 | 20.9 | 21.7 | 40.4 | 42.3 | 1.9 | 2.0 | *** |

All SHexCer species (*m/z* 806.5464, 820.5256, 822.5413) were spatially diffused across the upper hippocampus in *Sgsh*^R245H^, *Sgsh*^D31N^, and control mice, whereas SHex2Cer 36:1;O2 (*m/z* 968.5992) was broadly distributed in unaffected mice but strongly expressed in the CA1 hippocampal pyramidal cell layer in MPS IIIA mice (**Fig. 3**). Spatial distribution of *m/z* 865.5031 (likely BMP 44:12) appeared more localised to the hippocampal pyramidal cell layer and granule dentate gyrus (**Fig. 3**), distinguishing it from the alternate putative identity of isobaric phosphatidylglycerol (PG) 44:12. *m/z* 1179.7372, 1207.7685, 1382.8166, 1410.8479 (G_M3_ and G_M2_ ganglioside) MSI signals were clustered/punctate and localised to the hippocampal pyramidal cell layer in MPS IIIA mice, especially for the 36:1 species (**Fig. 3**). Conversely, the *m/z* 1470.8326 (putatively G_D3_ 36:1) MPS IIIA MSIs exhibited continuous, diffuse signals across the entire hippocampal section with strong intensity along the CA1 axis.

**Figure 3:**
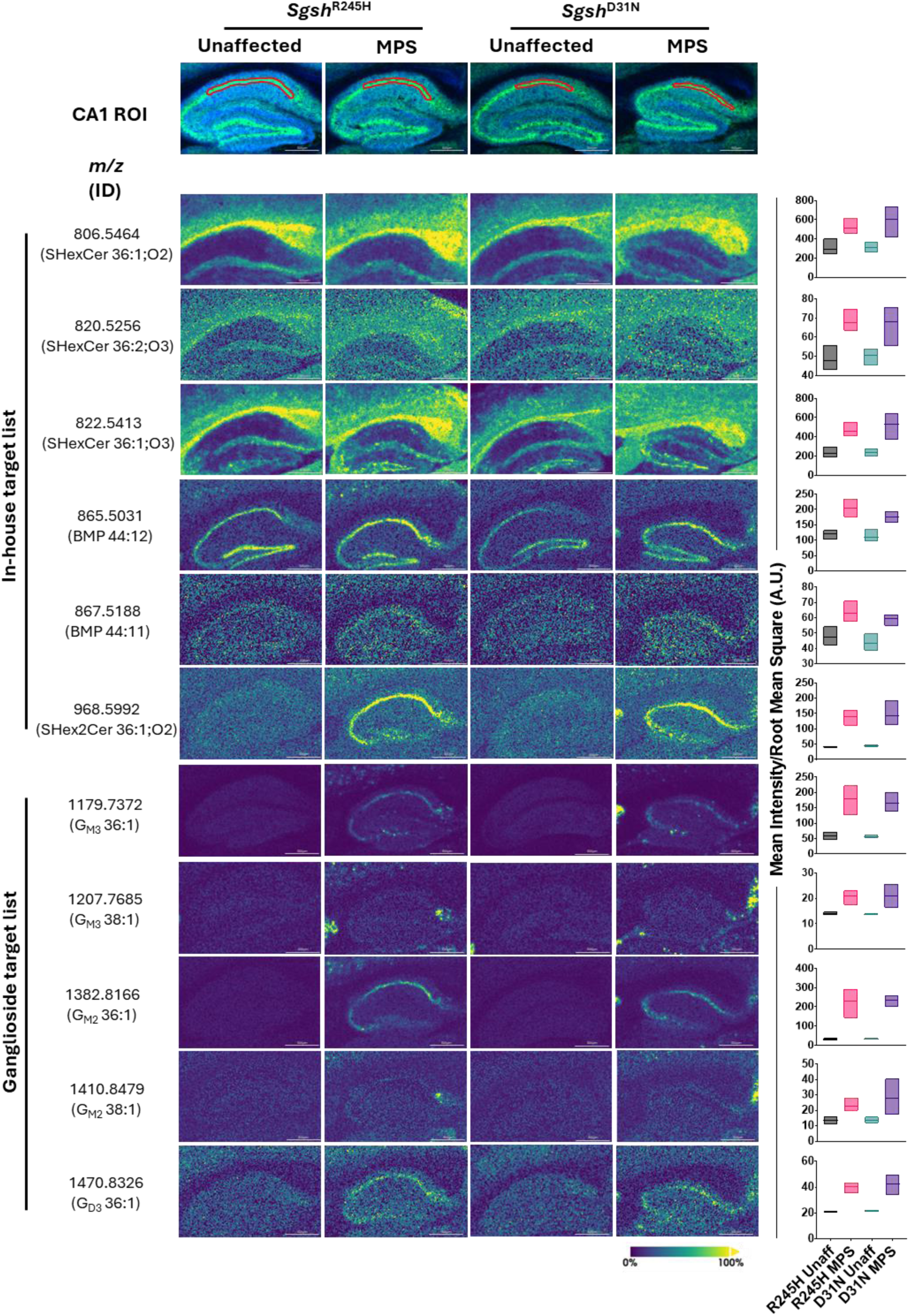
Representative mass spectrometry images from 20-week-old *Sgsh*^R245H^ and *Sgsh*^D31N^ MPS IIIA hippocampal sections. The CA1 hippocampus was defined as the region of interest (ROI). Curation against in-house and comprehensive ganglioside libraries identified 11 significant *m/z* lipid species. Signal intensities are displayed as ion images, with mean pixel intensity in the ROI normalised to root mean square and quantified in the far-right panels as box plots (min-max). Images are individual representatives of n=3 mice per group. BMP, bis(monoacylglycero)phosphate; G_M3_, monosialodihexosylganglioside; G_M2_, monosialotrihexosylganglioside; G_D3_, disialodihexosylganglioside; Unaff, unaffected; MPS, mucopolysaccharidosis type IIIA; AU, arbitrary units.

Lower abundant analytes such as SHexCer 36:2;O3, BMP 44:11, G_M3_ 38:1, G_M2_ 38:1, and G_D3_ 36:1, exhibited moderate changes in the MPS IIIA mice compared to controls (1.3-2.0-fold significantly increased) (**Table 1**). Similarly, the greatest abundance analytes, SHexCer 36:1;O2, SHexCer 36:2;O2, and BMP 44:12, were up to 2.2-fold significantly elevated in MPS IIIA mice. SHex2Cer 36:1;O2, G_M3_ 36:1, and G_M2_ 36:1 exhibited the greatest changes, wherein MPS IIIA mice exhibited 3.5-, 3.1-, 7.4- (*Sgsh*^R245H^), and 3.2-, 3.0-, 7.3- (*Sgsh*^D31N^) -fold increases compared to unaffected controls, respectively.

### MPS IIIA mice exhibit spatial memory deficits in the Morris Water Maze

We examined the behaviour of MPS IIIA mice using a test battery that evaluated memory and learning, motor impairment, and anxiety-like behaviours. All mice were naïve to the tests before commencing phenotyping at 22 weeks of age. The Morris Water Maze is a navigation task that measures spatial memory and learning. The escape latency improved with time in both genotypes, indicative of learning (**Supp**. **Fig 15a,b**). There were no significant genotype-specific differences apparent in either female or male mice. During the Probe phase, the platform was removed from the pool and memory retention was examined. Female MPS IIIA mice showed significantly reduced exploration of the target quadrant (expressed as a percentage of total exploration time) compared to similarly aged unaffected controls in the probe test (**Fig. 4a,b; Suppl. Fig. 15 c,d**). Likewise, male MPS IIIA mice exhibited a lower percentage of swim time and path length in the target quadrant compared to wild-type males. Unaffected males had greater accuracy in their search location and crossed the platform location more frequently than the MPS IIIA males (**Fig. 4c,d**). The swim speeds were significantly slower in MPS IIIA female mice but were comparable between unaffected and MPS IIIA males (**Suppl. Fig. 15e,f**).

**Figure 4:**
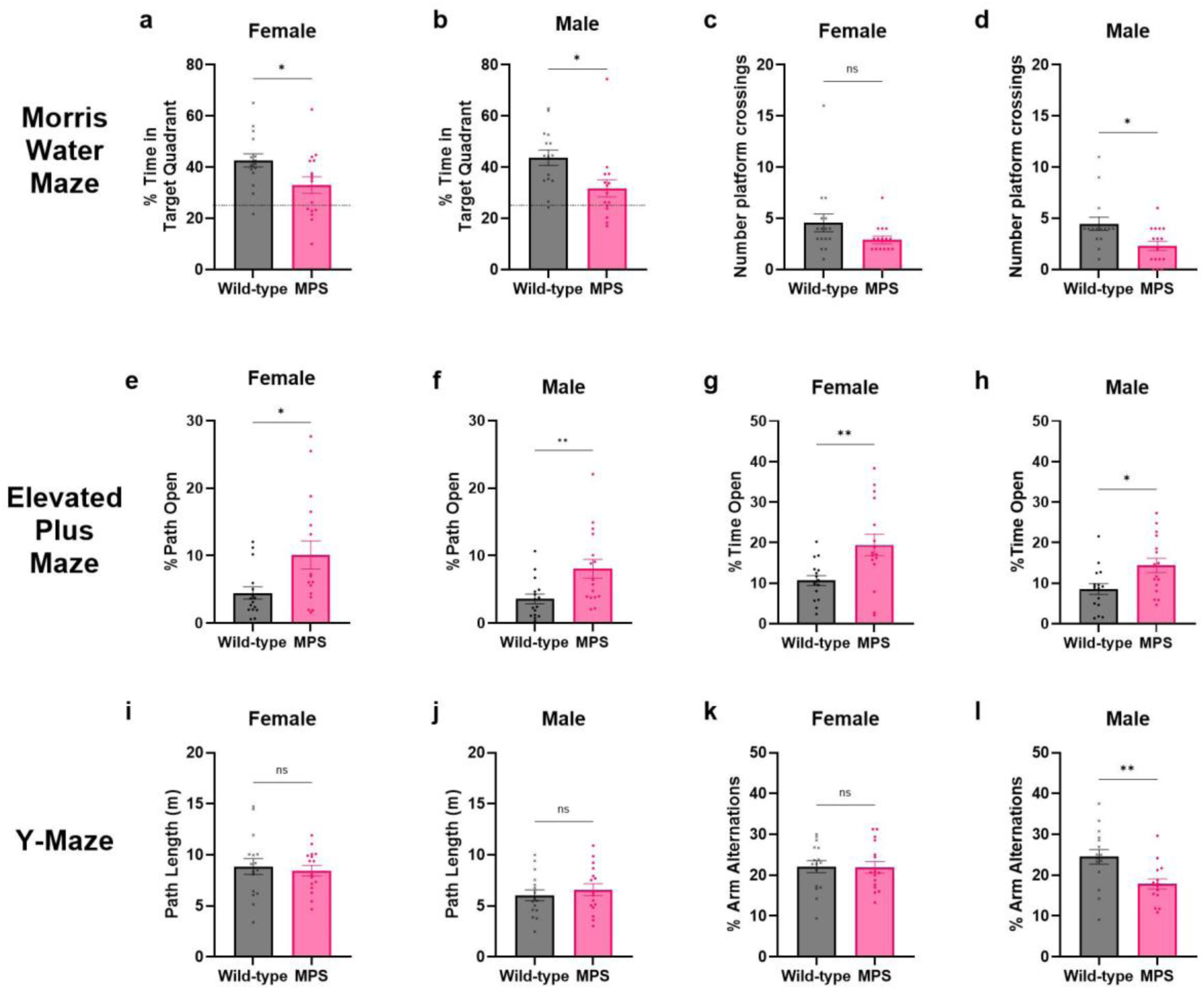
Behavioural phenotyping of MPS IIIA mice demonstrates the presence of memory and learning deficits and altered anxiety-related behaviours. Female and male wild-type and MPS IIIA mice were evaluated in a behavioural test battery. (a-d) In the Morris Water Maze probe test (22-23 weeks of age), the platform was removed from the pool and the exploration pattern was recorded. The dashed line indicates the 25% chance level if no preference is shown. The percentage of (a,b) time spent in the target quadrant and the (c,d) number of crossings of the location where the platform had previously been located was analysed with ANY-Maze software. (e-h) The exploration over 5 minutes in the Elevated Plus Maze was recorded at 24 weeks of age. Relative to wild-type mice, MPS IIIA mice displayed a greater percentage of (e,f) path length exploration as a percentage of total path length, and (g,h) the time spent exploring open arms as a percentage of total time spent in open or closed arms. (i-l) Working memory was examined using the Y-maze test at 25 weeks of age. (i,j) The distance travelled during exploration was not impacted by genotype. (k,l) The spontaneous arm alteration as a percentage of total arm entries revealed spatial short-term memory deficits in male MPS IIIA mice. n=15-16 mice/group; the data from one male wild-type mouse was excluded due to failing the visual test on day 7. ns, not significant; \**p*≤0.05, \*\**p*≤0.01.

### MPS IIIA mice show altered anxiety-related behaviours in the Elevated plus maze

The open field test assesses spontaneous locomotor activity and exploration of a novel arena. Compared to sex- and age-matched controls, MPS IIIA mice did not show any apparent differences in the total distance explored (path length), average walking speed nor the number of immobile episodes (**Supp. Fig. 15**). Following this, anxiety-like behaviours were examined in the Elevated Plus Maze. Spontaneous motor activity was unchanged in female and male MPS IIIA mice (path length and total arm entries; **Supp. Fig. 16a-d**). However, reduced anxiety responses were seen in MPS IIIA males, with more open arm entries, and greater time spent and path travelled on open arms (**Fig. 4. e-h; Supp. Fig. 16e,f**). Female MPS IIIA mice also showed reduced anxiety-related behaviours in the Elevated Plus Maze.

### Y-maze testing revealed spatial working memory impairment in male, but not female MPS IIIA mice

Short-term spatial working memory was assessed using the Y-maze test, which relies on the tendency of rodents to explore novel environments. Mice with intact working memory should recall which arms it has already visited and therefore exhibit a higher percentage of arm alternation. Similar exploratory activity was measured between unaffected and MPS IIIA mice for both sexes (path lengths; **Fig. 4i,j**). However, male wild-type mice showed a significantly higher percentage of arm alternations than male MPS IIIA mice (**Fig. 4l**). Spontaneous alternation was unchanged in female MPS IIIA mice (**Fig. 4k**).

### Gross motor function is unchanged in MPS IIIA mice

A series of motor tests that have previously shown differences in the naturally-occurring congenic *Sgsh*^D31N^ MPS IIIA mouse model were performed on transgenic *Sgsh*^R245H^ MPS IIIA mice. Age-matched wild-type and MPS IIIA mice had similar grip strength in the inverted grid test (time to fall), as well as latencies to turn in both the pole and negative geotaxis tests (**Supp. Fig. 17**). Hind-limb gait width and length was unchanged in MPS IIIA mice compared with wild-type mice (**Supp. Fig. 17**).

## Discussion

Here we describe the generation and characterisation of a novel transgenic MPS IIIA mouse model expressing the R245H *Sgsh* variant, an allele present in up to 58% of MPS IIIA patients (13). We confirmed that knock-in MPS IIIA mice exhibit significantly reduced SGSH enzymatic activity and progressively accumulate heparan sulfate in brain with age, validating the biochemical defect. We also demonstrate the age-dependent expansion of the endo-lysosomal compartment via LIMP2-immunostaining in *Sgsh*^R245H^ MPS IIIA mice. The time course of endo/lysosomal expansion is largely in agreement with that previously described in *Sgsh*^D31N^ and conditional *Sgsh* knockout MPS IIIA mice.

The accumulation of lysosomal substrate can trigger cellular dysfunction and pathological cascades that contribute to clinical disease. A robust neuroinflammatory response was found in *Sgsh*^R245H^ MPS IIIA mice, with increased numbers of isolectin B_4_-reactive, thus activated, microglia evident from as early as 3 weeks of age, akin to the early activated microglial response observed in *Sgsh*^D31N^ MPS IIIA (30) and knockout MPS IIIB (31) mice. These changes were accompanied by a progressive increase in the severity of astrogliosis in *Sgsh*^R245H^ MPS IIIA mouse brain, consistent with the timing and degree of GFAP staining found in *Sgsh*^D31N^ and conditional knockout MPS IIIA mouse models (5, 32). Widespread fibrillary gliosis has been observed in post-mortem brain tissue from siblings with MPS IIIB (33), highlighting the face validity of all three mouse models of MPS IIIA. That ‘clinically meaningful improvements in various neurobehavioral and quality-of-life outcomes’ were noted following provision of the IL-1beta receptor antagonist Anakinra to patients with MPS III (34) points to the key role neuroinflammation plays in MPS III.

Gangliosides G_M2_ and G_M3_ accumulate secondarily to heparan sulfate in the brain in MPS III and other cognate disorders (35). To further explore lipid perturbations in the *Sgsh*^R245H^ MPS IIIA brain, we have, for the first time for any MPS III disorder, employed a MALDI mass spectrometry-based imaging approach to both visualise and quantify lipid changes in diseased versus healthy mouse brain. We also compared the findings in *Sgsh*^R245H^ mice with those in the well-studied *Sgsh*^D31N^ mouse model, identifying 11 significantly altered species of common lipids in negative ion mode, all of which were elevated in both models (compared to unaffected controls), which indicates the presence of a similar pathological cascade and rate of disease progression in *Sgsh*^D31N^ and *Sgsh*^R245H^ MPS IIIA mice. As anticipated, the analysis revealed four significantly elevated *m/z* features, putatively assigned to G_M3_ and G_M2_ gangliosides, consistent with prior immunostaining outcomes and prior quantitation by LC-tandem mass spectrometry in *Sgsh*^D31N^ MPS IIIA mouse brain homogenates (5, 30, 36, 37). MALDI-MSI studies on cerebral cortex from knock-out mice with MPS II (Hunter syndrome) also demonstrated elevated G_M2_ and G_M3_ (38). In addition, however, we detected a further seven significantly altered *m/z* features not described in the MPS II mouse model; which may indicate more extensive lipid remodelling in brain associated with SGSH deficiency or may possibly reflect region-specific variations in lipid metabolism. We elected to study the hippocampus here, as it is critical for memory and learning behaviours (29), particularly spatial and episodic memory processing, in which both *Sgsh*^R245H^ (here) and *Sgsh*^D31N^mice (4) are deficient. G_M3_ and G_M2_ MSI signals were clustered/punctate in MPS IIIA mice (**Fig. 3**), consistent with their secondary lysosomal accumulation, which has been attributed to heparan sulfate-induced inhibition of lysosomal hydrolases, sialidase (neuraminidase 1) and β-hexosaminidase (39).

It was also unsurprising that the 36:1 and 38:1 species were implicated given that the likely corresponding d18:1/18:0 and d18:1/20:0 species reflect typical ganglioside compositions in the brain (40) which are elevated in the other MPS IIIA mouse models (5, 36, 41, 42). Whilst secondary G_D3_ accumulation has been reported in the MPS I mouse (43) and the MPS IIID goat (44), altered G_D3_ concentrations are seldom described in MPS disorders, with only one other report noting increased levels in the brain stem of the *Sgsh*^D31N^ MPS IIIA mouse at 1 month, but not 6 months of age. There were no changes in G_D3_ seen in the cerebellum, cortex, or sub-cortex (42). Given the broad spatial distribution, and the previous association of G_D3_ with astrocyte activation (45), G_D3_ elevation may reflect increased reactive astrogliosis, consistent with elevated GFAP staining in the inferior colliculus and cerebral rostral cortex in the *Sgsh*^R245H^ MPS IIIA mice. The broad distribution may further suggest increased G_D3_ synthesis rather than impaired lysosomal turnover, however, further work is required to confirm this postulate.

The other six *m/z* whose levels were significantly increased in the MPS IIIA mouse CA1 hippocampal region (c.f. unaffected mice) were putatively identified as BMP 44:12 (*m/z* 865.5031), BMP 44:11 (*m/z* 867.5188), SHexCer 36:1;O2 (*m/*z 806.5464), SHexCer 36:2;O3 (*m/z* 820.5256), SHexCer 36:1;O3 (*m/z* 822.5413), and SHex2Cer 36:1;O2 (*m/z* 968.5992) (**Fig. 3**; **Table 1**). Increased BMP has been reported in MPS IIIA mouse brain previously (41, 46), however, the mechanistic cause remains unknown. Reduced BMP catabolism might be a result of impaired endo-lysosomal clearance or altered synthesis. Drug-induced upregulation of BMP synthesis and/or direct supplementation of BMP *in vitro* has been shown to reduce lipid-associated pathologies in other neurodegenerative disorders such as Niemann-Pick C and frontotemporal dementia, stimulating lipid metabolism/egress and reducing lysosomal storage material (47, 48). In fact, lipid remodelling and other compensatory actions are the subject of increasing interest in the lysosomal storage disorder field (49–51) (**Fig. 2b**). For example, BMP has been shown to activate GM2 activator protein and hexosaminidase A *in vitro* (52), which could suggest that the increase in BMP is a compensatory mechanism designed to offset the secondary accumulations of G_M2_. Further clarification of BMP’s role in MPS IIIA is warranted.

SHexCer (also known as sulfatide) is highly enriched in the myelin sheath, serving as the major glycosphingolipid component of oligodendrocytes (53) and induces cellular toxicity, ROS production and mitochondrial stress (54). Elevated SHexCer in the CA1 hippocampal region is largely atypical of neurodegenerative disease, though an increase in SHexCer (d36:1; *m/*z 806.545) was reported in the corpus callosum of the Shiverer mouse model of demyelination using MALDI-MSI (55), accompanied by a decrease in SHexCer (d42:2; *m/z* 888.623). Most neurodegenerative disorders with white matter and/or dysmyelination pathologies exhibit reduced levels of sulfatide in the brain, including across predominantly grey matter structures such as the cortex, hippocampus, and cerebellum, with a general decrease as the brain ages (56–58). The reductions observed in white matter regions in other disorders might reflect dysmyelination or oligodendrocyte loss, whereas the elevated grey matter sulfatide levels observed here may instead reflect altered lipid metabolism or turnover. Interestingly, a recent MALDI-MSI analysis of brain from mice with Gracile Axonal Dystrophy showed elevated concentrations of d18:1/18:0 and d18:1/18:0-OH sulfatides in the dorsomedial medulla, suggesting a more critical role for sulfatides in axonal regulation (59). Interestingly, only SHexCer comprising 36 carbons was significantly altered in the MPS IIIA mice compared to controls, most probably corresponding to the d18:1 sphingosine backbone with varying levels of acyl chain saturation and hydroxylation. A prior report in MPS IIIA mice has shown decreased concentrations of d18:1/24:1 sulfatide in the brain stem at 6 months, but no changes were observed in the cerebellum, cortex, or sub-cortex; d18:1/18:0 sulfatide was also unchanged (42). Given the importance of 36:1 non-hydroxylated and hydroxylated SHexCer in early oligodendrocyte maturation (60), and depending on the age of onset of this abnormality, the increased concentrations of 36:1 sulfatides might indicate prolonged oligodendrocyte immaturity in MPS IIIA.

MPS IIIA mice also exhibited profound elevations of *m/z* 968.5992, the sulfated-glycosphingolipid SHex2Cer 36:1;O2 in the CA1 field of hippocampus. Unlike canonical sulfatides derived from galactosylceramide, SHex2Cer is generated via direct sulfation of lactosylceramide and therefore arises from a key branch point in glycosphingolipid metabolism, shared with G_M3_ and other downstream gangliosides (61). Consistent with this, sphingolipid remodelling accompanied by prominent SHex2Cer accumulation has been reported in *Galgt1/Siat9* double-null mice, in which ganglioside synthesis is blocked (62). In the context of substantial G_M3_, G_M2_, and G_D3_ accumulation in MPS IIIA mouse brain here, elevated SHex2Cer may reflect altered metabolic flux through lactosylceramide, potentially arising from dysregulated glycosphingolipid processing. Alternatively, impaired hydrolytic capacity of arylsulfatase A could also explain the elevated levels of SHexCer and SHex2Cer in MPS IIIA mouse brain. However, given that SHex2Cer is most typically found in visceral tissues (63), the substantially elevated levels seen here more likely reflects changes in its anabolism, rather than catabolism. Metabolic tracing studies are required to clarify pathway dynamics and determine the contribution of sulfatide dysregulation to MPS IIIA pathology.

Consequent to the metabolic and histopathological changes observed in the brain, MPS IIIA male and female mice exhibited a reduced spatial bias towards the quadrant where the escape platform had previously been located compared with sex-matched controls in the Morris Water Maze probe test. This suggests that MPS IIIA mice display a reduced ability to remember the platform location, indicative of spatial memory deficits, albeit swim speed was slightly slower in MPS IIIA females and so a motor phenotype component cannot be excluded. Short-term working memory deficits were also displayed by male MPS IIIA mice in the Y-maze (reduced percentage arm alternation). The overall exploration of the maze did not influence this parameter as the total distance walked was not impacted by genotype. Likewise, male and female MPS IIIA mice displayed reduced anxiety-like behaviours during their exploration of the open arms in the Elevated Plus Maze without an overall change to gross exploration (total arm entries; path length). These findings are consistent with those described in the *Sgsh*^D31N^ mouse model (25, 26), and together with the similar lipidomic outcomes described above, indicate that both the D31N and R245H mutations in *Sgsh* cause rapidly progressing brain disease when modelled in mice.

Under this testing protocol, which evaluated behavioural responses as part of a test battery, the overall open-field locomotor activity was unaltered in *Sgsh*^R245H^ MPS IIIA mice, and there was also no evidence of motor deficits. This contrasts with the open field activity changes and motor phenotype that has been frequently documented in *Sgsh*^D31N^ MPS IIIA mice (25, 26, 28, 64). We have previously found that repeated handling during clinical monitoring and daily treatments in *Sgsh*^D31N^ mice affected exploration, preventing us from measuring differences in open-field locomotor activity between genotypes (65). The order of the behavioural tests can also confound testing outcomes in the later tests, potentially due to the cumulative stress of handling and/or familiarisation with experimenter handling. Training history plays a significant role in how the mice perform in subsequent tests. McIlwain *et al*. (66) evaluated male C57BL/6J mice in a test battery ordered in what was thought to be the least to most invasive. Naïve-tested mice were more active in an open field arena than those that were evaluated in a neurological screen prior to open field testing (66). Battery-tested mice also had improved motor coordination on the rotorod compared to the performance of naïve-tested mice (66). Evaluating the same mouse over multiple assays allows correlations to be made between several behavioural measures and reduces the overall number of mice required, which is advantageous from an ethical perspective. However, careful consideration should be given to the order and number of tests in a battery. Re-evaluation of the locomotor activity and motor phenotype in naïve-cohorts of *Sgsh*^R245H^ mice would be valuable to address the effects of testing order.

Several transgenic MPS IIIA mouse models have been developed to examine disease etiology, evaluate therapies or to serve as research tools. For example, at least two immune-deficient MPS IIIA models have been established where affected MPS IIIA mice lacked Rag2, CD47 and Il2rg genes (9), or were interbred with NOD/SCID/GammaC chain null mice (67). These models allow the xenotransplantation and engraftment of human neural stem cells or iPSC-derived neural progenitor cells. MPS IIIA mice with ubiquitous expression of a GFP reporter in peripheral blood leukocytes have also been generated, allowing identification of donor cells in transplantation models (68). Furthermore, two independent conditional knockout *Sgsh* MPS IIIA mouse models have been created. In the MPS IIIA mouse model developed by Dwyer et al. (6), study of the impact of cell-autonomous *Sgsh* deletion in several neural cell types was hampered by apparent cross-correction of *Sgsh-*deleted cells by unaltered cells. The other conditional knockout MPS IIIA model has ubiquitous *Sgsh* knockout (5). Side-by-side comparison of conditional knockout MPS IIIA mice with *Sgsh*^D31N^ mice expressing residual SGSH enzyme activity of ∼3% wild-type levels (5) demonstrated that the onset of cognitive and motor behavioural changes was slightly earlier in the conditional knockout MPS IIIA mice despite both MPS IIIA models accumulating heparan sulfate and G_M2_/G_M3_ glycosphingolipids at similar rates, suggesting that these storage products were not the primary determinant for symptom generation (5).

The first and thus most widely used MPS IIIA mouse model is the naturally-occurring *Sgsh*^D31N^ MPS IIIA model that was originally identified in mice with a mixed genetic background (3), and was later back- crossed onto the C57BL/6 background by our laboratory (4) and Jackson Laboratories. Its validity and suitability as an authentic MPS IIIA animal model are highlighted by the many clinical trials that have emanated from testing in these mice. However, despite the clear relevance of the *Sgsh*^D31N^ model, this mutation has not been described in MPS IIIA patients to our knowledge. This is particularly pertinent to the development of therapeutics, such as pharmacological chaperones, that require validation across different mutant proteins. Further, the outcome of recent studies in MPS IIIC mice (17) led to a postulate that there is a toxic gain-of-function resulting from the presence of misfolded mutant lysosomal enzyme(s), with missense mutants displaying earlier decline in neurological function than knockout mice. The availability of the *Sgsh*^R245H^ mouse model will allow this hypothesis to be further explored in another MPS III subtype.

## Conclusions

*Sgsh*^R245H^ MPS IIIA mice exhibit phenotypic, biochemical and neuropathological changes that are similar in age-of-onset and rate of progression to other previously described MPS IIIA mutant mouse lines, and importantly, the new model also exhibits face-validity to the human disorder, thus validating the usefulness of this mouse model in studying MPS IIIA disease pathogenesis, biomarker validation and evaluation of innovative therapeutic approaches. MALDI-MSI studies facilitated detection, imaging and quantitation of elevated levels of novel putative sulfatide substrates in the *Sgsh*^R245H^ and *Sgsh*^D31N^ mouse model hippocampal CA1 field, with further studies required to determine the mechanism leading to, and implication of, their presence.

## Acknowledgements

This study was funded by the Sanfilippo Children’s Foundation and Fondation Sanfilippo Suisse in a grant to KMH and AAL. The authors acknowledge and thank the South Australian Genome Editing (SAGE) Facility, who are supported by Phenomics Australia and are funded by the Australian Government through the National Collaborative Research Infrastructure Strategy (NCRIS) program. We also acknowledge and thank the facilities, and the scientific and technical assistance of Microscopy Australia (ROR: 042mm0k03) enabled by NCRIS and the government of South Australia at Flinders Microscopy and Microanalysis (ROR: 04z91ja70), Flinders University (ROR: 01kpzv902). The graphical abstract was created in BioRender. Bowen, A. (2027) https://BioRender.com/9ngl17t. We thank Dr Stuart Read (SAHMRI) for the use of the Y-maze apparatus, and Associate Professor Arne Ittner (Flinders University) for access to the behavioural analysis software.

## Data availability statement

The data from this study are available upon reasonable request.

## Ethics approval statement

All animal studies were approved by the Institutional Animal Ethics Committee in accordance with the 2013 National Health and Medical Research Council’s Australian Code For The Care And Use Of Animals for Scientific Purposes, 8^th^ edition (approval number 5726).

