## Supplementary Information for "Cognitive impairment and progressive neuroinflammation in mucopolysaccharidosis IIIA mice expressing the R245H Sgsh variant"

**Data availability statement:** The data from this study are available upon reasonable request.

**Funding statement:** We thank the Sanfilippo Children's Foundation (Australia) and Fondation Sanfilippo Suisse (Switzerland) for funding this study.

**Conflict of interest disclosure:** All authors declare that they have no conflict of interest.

**Ethics approval statement**: All animal studies were approved by the Institutional Animal Ethics Committee in accordance with the 2013 National Health and Medical Research Council's Australian Code For The Care And Use Of Animals for Scientific Purposes, 8th edition (approval number 5726).

### Supplementary Methods

#### ***Behavioural testing***

Morris water maze [cohort 1, 22-weeks-old; cohort 2, 23-weeks-old; (1)]. Mice were tested in a 1.2-m diameter pool filled with 20-23°C water made opaque with non-toxic white paint (Spring) containing a round, clear plastic, 10-cm diameter platform positioned 1-cm below the level of the water. Two monochromatic visual poster cues were located on the walls adjacent to the platform. Acquisition phase (days 1-5): the time taken and the distance swum to find the hidden platform was recorded. There were four trials daily starting at each of the quadrants. Mice were recovered on heating pads between trials (>10 min inter-trial interval). If the mouse failed to find the platform within 90s, it was guided there and allowed ~10s to learn the platform position before removal from the pool. Probe phase (day 6): the platform was removed from the pool and mice were assessed in one 90s test starting in the quadrant opposite where the platform was previously located. Visual test (day 7). A colourful object was placed on the platform, which was in a new quadrant, and the visual cues were removed. Mice were tested up to 4 times for a maximum of 90 s, or until a latency of less than 10 seconds was achieved.

Open field activity [24-weeks-old (2)]. Mice were placed into the corner of a 30x50 cm container facing the wall and exploration was recorded for 5 min.

Elevated plus maze [24-weeks-old (2)]. The apparatus consisted of a "+"-shaped maze with arms 7-cm wide x 50-cm long. It was elevated 50-cm above the floor. Two arms were enclosed with black walls 40-cm high ("closed" arms), and two had no walls ("open arms"). The floor was white. Mice were placed in the central square facing an open arm at the start of the test, and exploration over 5-min was recorded.

Y-maze test [25-weeks-old (3)]. The apparatus was constructed from grey plastic (3-cm bottom and 10-cm top width, 40-cm long and 12-cm high radiating out from the central triangle. The maze was positioned 60-cm above the floor. Mice were placed in the centre of the Y-maze and allowed to explore for 5 minutes. The arm that the mouse facing at the start of the trial was alternated.

Motor tests [25-weeks-old (4, 5)]. Neuromuscular grip strength was measured by placing mice on a wire grid (15x30 cm) and inverting it horizontally above a box filled with shredded paper. The time until the mouse fell from the grid was measured up to 60s. A fail was recorded if the mouse fell within 60 s. Pole test: Mice were placed facing head-upward on the top of a vertical rough-surfaced pole (diameter 8-mm, height 55-cm) and the time taken to orient down and climb down to the home cage was measured up to a maximum of 120s. If the mouse fell during the test, a 'fall' was recorded and the mouse was given a total score of 120 s. Negative geotaxis: Mice were placed on a wire grid (15 x 30 cm) and the grid was upended so that the mouse faced towards the floor above a box filled with shredded paper. The time taken for the mouse to re-orient and face the ceiling up to a maximum of 30 s was measured. A fail was recorded if the mouse fell within 30 s. Gait: Mice had their hind paws dipped in food colouring and walked freely down a corridor lined with paper towards a dark goalbox. The length and width between the footprints were measured from at least three strides and two replicate tracks.

| Target | Titration | Antibody | Manufacturer and Catalogue Number |
|---|---|---|---|
| **Lysosomal integral membrane protein-2** | 1:500<br><br>1:2000 | Monoclonal mouse anti-LIMP2<br><br>Biotinylated donkey anti-mouse IgG | In-house (6)<br><br>Jackson ImmunoResearch; #715-065-150 |
| **Glial fibrillary acidic protein** | 1:13,000<br><br>1:2000 | Polyclonal rabbit anti-GFAP<br><br>Biotinylated donkey anti-rabbit IgG | Agilent (DAKO); #Z033401-2<br><br>Jackson ImmunoResearch; #711-065-152 |
| **Isolectin $B_4$** | 1:100 | Peroxidase conjugated, isolectin $B_4$ from *Bandeiraea simplicifolia* (*Griffonia simplicifolia*) | Sigma-Aldrich; #L5391 |

**Supplementary Table 1. Details of antibodies and lectin reagents.**

| *m/z* | *Tentative Identity* | *Ion Mode* |
|---|---|---|
| *407.2210* | LPA 16:1 | $[M-H]^-$ |
| *409.2367* | LPA 16:0 | $[M-H]^-$ |
| *420.2277* | LPA 17:1 | $[M-H]^-$ |
| *422.2433* | LPA 17:0 | $[M-H]^-$ |
| *433.2367* | LPA 18:2 | $[M-H]^-$ |
| *435.2523* | LPA 18:1 | $[M-H]^-$ |
| *437.2680* | LPA 18:0 | $[M-H]^-$ |
| *450.2746* | LPA 19:0 | $[M-H]^-$ |
| *452.2789* | LPE 16:0 | $[M-H]^-$ |
| *457.2367* | LPA 20:4 | $[M-H]^-$ |
| *461.2680* | LPA 20:2 | $[M-H]^-$ |
| *463.2836* | LPA 20:1 | $[M-H]^-$ |
| *465.2993* | LPA 20:0 | $[M-H]^-$ |
| *465.3039* | Cholesterol Sulfate | $[M-H]^-$ |
| *478.2945* | LPE 18:1 | $[M-H]^-$ |
| *480.3102* | LPE 18:0 | $[M-H]^-$ |
| *481.2367* | LPA 22:6 | $[M-H]^-$ |
| *483.2523* | LPA 22:5 | $[M-H]^-$ |
| *483.2734* | LPG 16:0 | $[M-H]^-$ |
| *485.2680* | LPA 22:4 | $[M-H]^-$ |
| *497.2880* | LPG 17:0 | $[M-H]^-$ |
| *500.2789* | LPE 20:4 | $[M-H]^-$ |
| *506.3258* | LPE 20:1 | $[M-H]^-$ |
| *507.2734* | LPG 18:2 | $[M-H]^-$ |
| *509.2891* | LPG 18:1 | $[M-H]^-$ |
| *511.3047* | LPG 18:0 | $[M-H]^-$ |
| *522.2843* | LPS 18:1 | $[M-H]^-$ |
| *524.2789* | LPE 22:6 | $[M-H]^-$ |
| *555.2734* | LPG 22:6 | $[M-H]^-$ |
| *568.2687* | LPS 22:6 | $[M-H]^-$ |
| *569.2738* | LPI 16:1 | $[M-H]^-$ |
| *570.2843* | LPS 22:5 | $[M-H]^-$ |
| *571.2895* | LPI 16:0 | $[M-H]^-$ |
| *597.3051* | LPI 18:1 | $[M-H]^-$ |
| *599.3208* | LPI 18:0 | $[M-H]^-$ |
| *619.2895* | LPI 20:4 | $[M-H]^-$ |
| *645.4507* | PA 32:1 | $[M-H]^-$ |
| *647.4663* | PA 32:0 | $[M-H]^-$ |
| *662.4772* | PE 30:0 | $[M-H]^-$ |
| *670.5269* | HexCer 32:1 | $[M-H]^-$ |
| *671.4663* | PA 34:2 | $[M-H]^-$ |

| | | |
|---|---|---|
| *673.4820* | PA 34:1 | $[M-H]^-$ |
| *675.4976* | PA 34:0 | $[M-H]^-$ |
| *676.5293* | PE O-32:0 | $[M-H]^-$ |
| *687.4965* | PA 35:1 | $[M-H]^-$ |
| *688.4929* | PE 32:1 | $[M-H]^-$ |
| *689.5121* | PA 35:0 | $[M-H]^-$ |
| *690.5085* | PE 32:0 | $[M-H]^-$ |
| *693.4507* | PA 36:5 | $[M-H]^-$ |
| *695.4663* | PA 36:4 | $[M-H]^-$ |
| *696.5426* | HexCer 34:2 | $[M-H]^-$ |
| *697.4820* | PA 36:3 | $[M-H]^-$ |
| *698.5582* | HexCer 34:1 | $[M-H]^-$ |
| *699.4976* | PA 36:2 | $[M-H]^-$ |
| *700.5293* | PE O-34:2 / PE P-34:1 | $[M-H]^-$ |
| *701.5133* | PA 36:1 | $[M-H]^-$ |
| *702.5449* | PE O-34:1 / PE P-34:0 | $[M-H]^-$ |
| *703.5289* | PA 36:0 | $[M-H]^-$ |
| *709.4808* | PA 37:4 | $[M-H]^-$ |
| *710.5582* | HexCer 35:2 | $[M-H]^-$ |
| *712.4929* | PE 34:3 | $[M-H]^-$ |
| *713.5121* | PA 37:2 | $[M-H]^-$ |
| *714.5085* | PE 34:2 | $[M-H]^-$ |
| *715.5278* | PA 37:1 | $[M-H]^-$ |
| *716.5242* | PE 34:1 | $[M-H]^-$ |
| *717.4507* | PA 38:7 | $[M-H]^-$ |
| *718.5398* | PE 34:0 | $[M-H]^-$ |
| *719.4663* | PA 38:6 | $[M-H]^-$ |
| *720.4604* | PE 35:6 | $[M-H]^-$ |
| *720.5426* | HexCer 36:4 | $[M-H]^-$ |
| *721.4820* | PA 38:5 | $[M-H]^-$ |
| *722.4761* | PE 35:5 | $[M-H]^-$ |
| *722.5136* | PE O-36:5 / PE P-36:4 | $[M-H]^-$ |
| *723.4976* | PA 38:4 | $[M-H]^-$ |
| *724.4917* | PE 35:4 | $[M-H]^-$ |
| *724.5293* | PE O-36:4 / PE P-36:3 | $[M-H]^-$ |
| *725.5133* | PA 38:3 | $[M-H]^-$ |
| *726.5074* | PE 35:3 | $[M-H]^-$ |
| *726.5449* | PE O-36:3 / PE P-36:2 | $[M-H]^-$ |
| *726.5895* | HexCer 36:1 | $[M-H]^-$ |
| *727.5289* | PA 38:2 | $[M-H]^-$ |
| *728.5230* | PE 35:2 | $[M-H]^-$ |
| *728.5606* | PE O-36:2 / PE P-36:1 | $[M-H]^-$ |

| | | |
|---|---|---|
| *729.5446* | PA 38:1 | $[M-H]^-$ |
| *730.5387* | PE 35:1 | $[M-H]^-$ |
| *730.5762* | PE O-36:1 / PE P-36:0 | $[M-H]^-$ |
| *733.4808* | PA 39:6 | $[M-H]^-$ |
| *734.4772* | PE 36:6 | $[M-H]^-$ |
| *734.4984* | PS 32:0 | $[M-H]^-$ |
| *735.4965* | PA 39:5 | $[M-H]^-$ |
| *736.4929* | PE 36:5 | $[M-H]^-$ |
| *737.5121* | PA 39:4 | $[M-H]^-$ |
| *738.5085* | PE 36:4 | $[M-H]^-$ |
| *740.5242* | PE 36:3 | $[M-H]^-$ |
| *742.5398* | PE 36:2 | $[M-H]^-$ |
| *743.4663* | PA 40:8 | $[M-H]^-$ |
| *744.5555* | PE 36:1 | $[M-H]^-$ |
| *745.4820* | PA 40:7 | $[M-H]^-$ |
| *746.5136* | PE O-38:7 / PE P-38:6 | $[M-H]^-$ |
| *746.5711* | PE 36:0 | $[M-H]^-$ |
| *747.4976* | PA 40:6 | $[M-H]^-$ |
| *748.4917* | PE 37:6 | $[M-H]^-$ |
| *748.5293* | PE O-38:6 / PE P-38:5 | $[M-H]^-$ |
| *749.4253* | PI 28:2 | $[M-H]^-$ |
| *749.5133* | PA 40:5 | $[M-H]^-$ |
| *749.5344* | PG 34:0 | $[M-H]^-$ |
| *750.5074* | PE 37:5 | $[M-H]^-$ |
| *750.5449* | PE O-38:5 / PE P-38:4 | $[M-H]^-$ |
| *751.4409* | PI 28:1 | $[M-H]^-$ |
| *751.4550* | PG 35:6 | $[M-H]^-$ |
| *751.5289* | PA 40:4 | $[M-H]^-$ |
| *752.5230* | PE 37:4 | $[M-H]^-$ |
| *752.5606* | PE O-38:4 / PE P-38:3 | $[M-H]^-$ |
| *753.4566* | PI 28:0 | $[M-H]^-$ |
| *753.5446* | PA 40:3 | $[M-H]^-$ |
| *754.5387* | PE 37:3 | $[M-H]^-$ |
| *754.5762* | PE O-38:3 / PE P-38:2 | $[M-H]^-$ |
| *756.5919* | PE O-38:2 / PE P-38:1 | $[M-H]^-$ |
| *758.4984* | PS 34:2 | $[M-H]^-$ |
| *760.4929* | PE 38:7 | $[M-H]^-$ |
| *760.5140* | PS 34:1 | $[M-H]^-$ |
| *762.5085* | PE 38:6 | $[M-H]^-$ |
| *764.5242* | PE 38:5 | $[M-H]^-$ |
| *765.4718* | PG 36:6 | $[M-H]^-$ |
| *766.5398* | PE 38:4 | $[M-H]^-$ |

| | | |
|---|---|---|
| *767.4663* | PA 42:10 | $[M-H]^-$ |
| *768.4816* | PS 35:4 | $[M-H]^-$ |
| *768.5555* | PE 38:3 | $[M-H]^-$ |
| *769.4820* | PA 42:9 | $[M-H]^-$ |
| *769.5031* | PG 36:4 | $[M-H]^-$ |
| *770.4972* | PS 35:3 | $[M-H]^-$ |
| *770.5136* | PE O-40:9 / PE P-40:8 | $[M-H]^-$ |
| *770.5711* | PE 38:2 | $[M-H]^-$ |
| *771.4976* | PA 42:8 | $[M-H]^-$ |
| *771.5188* | PG 36:3 | $[M-H]^-$ |
| *772.5129* | PS 35:2 | $[M-H]^-$ |
| *772.5293* | PE O-40:8 / PE P-40:7 | $[M-H]^-$ |
| *772.5868* | PE 38:1 | $[M-H]^-$ |
| *773.5133* | PA 42:7 | $[M-H]^-$ |
| *773.5344* | PG 36:2 | $[M-H]^-$ |
| *774.5285* | PS 35:1 | $[M-H]^-$ |
| *774.5449* | PE O-40:7 / PE P-40:6 | $[M-H]^-$ |
| *775.5501* | PG 36:1 | $[M-H]^-$ |
| *776.4994* | SHexCer 34:2 | $[M-H]^-$ |
| *776.5230* | PE 39:6 | $[M-H]^-$ |
| *776.5442* | PS 35:0 | $[M-H]^-$ |
| *776.5606* | PE O-40:6 / PE P-40:5 | $[M-H]^-$ |
| *777.5446* | PA 42:5 | $[M-H]^-$ |
| *777.5657* | PG 36:0 | $[M-H]^-$ |
| *778.4671* | PS 36:6 | $[M-H]^-$ |
| *778.5151* | SHexCer 34:1 | $[M-H]^-$ |
| *778.5762* | PE O-40:5 / PE P-40:4 | $[M-H]^-$ |
| *779.4722* | PI 30:1 | $[M-H]^-$ |
| *780.4827* | PS 36:5 | $[M-H]^-$ |
| *780.5919* | PE O-40:4 / PE P-40:3 | $[M-H]^-$ |
| *781.4879* | PI 30:0 | $[M-H]^-$ |
| *782.4772* | PE 40:10 | $[M-H]^-$ |
| *782.4984* | PS 36:4 | $[M-H]^-$ |
| *784.5140* | PS 36:3 | $[M-H]^-$ |
| *786.5085* | PE 40:8 | $[M-H]^-$ |
| *786.5297* | PS 36:2 | $[M-H]^-$ |
| *787.5489* | PG 37:2 | $[M-H]^-$ |
| *788.5242* | PE 40:7 | $[M-H]^-$ |
| *788.5453* | PS 36:1 | $[M-H]^-$ |
| *790.5398* | PE 40:6 | $[M-H]^-$ |
| *791.4663* | PA 44:12 | $[M-H]^-$ |
| *792.4816* | PS 37:6 | $[M-H]^-$ |

| | | |
|---|---|---|
| *792.4943* | SHexCer 34:2 | $[M-H]^-$ |
| *792.5555* | PE 40:5 | $[M-H]^-$ |
| *793.4820* | PA 44:11 | $[M-H]^-$ |
| *793.5031* | PG 38:6 | $[M-H]^-$ |
| *794.4972* | PS 37:5 | $[M-H]^-$ |
| *794.5100* | SHexCer 34:1 | $[M-H]^-$ |
| *794.5136* | PE O-42:11 / PE P-42:10 | $[M-H]^-$ |
| *794.5711* | PE 40:4 | $[M-H]^-$ |
| *795.4976* | PA 44:10 | $[M-H]^-$ |
| *795.5188* | PG 38:5 | $[M-H]^-$ |
| *796.5129* | PS 37:4 | $[M-H]^-$ |
| *796.5256* | SHexCer 34:0 | $[M-H]^-$ |
| *796.5293* | PE O-42:10 / PE P-42:9 | $[M-H]^-$ |
| *796.5868* | PE 40:3 | $[M-H]^-$ |
| *797.5133* | PA 44:9 | $[M-H]^-$ |
| *797.5344* | PG 38:4 | $[M-H]^-$ |
| *798.5285* | PS 37:3 | $[M-H]^-$ |
| *802.5762* | PE O-42:7 / PE P-42:6 | $[M-H]^-$ |
| *803.4722* | PI 32:3 | $[M-H]^-$ |
| *804.4827* | PS 38:7 | $[M-H]^-$ |
| *804.5307* | SHexCer 36:2 | $[M-H]^-$ |
| *805.4879* | PI 32:2 | $[M-H]^-$ |
| *806.4984* | PS 38:6 | $[M-H]^-$ |
| *806.5464* | SHexCer 36:1 | $[M-H]^-$ |
| *807.5035* | PI 32:1 | $[M-H]^-$ |
| *807.5176* | PG 39:6 | $[M-H]^-$ |
| *808.4929* | PE 42:11 | $[M-H]^-$ |
| *808.5140* | PS 38:5 | $[M-H]^-$ |
| *808.6678* | HexCer 42:2 | $[M-H]^-$ |
| *809.5192* | PI 32:0 | $[M-H]^-$ |
| *809.5333* | PG 39:5 | $[M-H]^-$ |
| *810.5085* | PE 42:10 | $[M-H]^-$ |
| *810.5297* | PS 38:4 | $[M-H]^-$ |
| *810.6834* | HexCer 42:1 | $[M-H]^-$ |
| *812.5242* | PE 42:9 | $[M-H]^-$ |
| *812.5453* | PS 38:3 | $[M-H]^-$ |
| *812.6545* | PE O-42:2 / PE P-42:1 | $[M-H]^-$ |
| *815.4123* | DLCL 28:2 | $[M-H]^-$ |
| *816.5555* | PE 42:7 | $[M-H]^-$ |
| *816.5766* | PS 38:1 | $[M-H]^-$ |
| *818.5711* | PE 42:6 | $[M-H]^-$ |
| *819.5188* | PG 40:7 | $[M-H]^-$ |

| | | |
|---|---|---|
| *820.5129* | PS 39:6 | $[M-H]^-$ |
| *820.5256* | SHexCer 36:2 | $[M-H]^-$ |
| *820.5293* | PE O-44:12 / PE P-44:11 | $[M-H]^-$ |
| *821.5344* | PG 40:6 | $[M-H]^-$ |
| *822.5285* | PS 39:5 | $[M-H]^-$ |
| *822.5413* | SHexCer 36:1 | $[M-H]^-$ |
| *822.5449* | PE O-44:11 / PE P-44:10 | $[M-H]^-$ |
| *823.5501* | PG 40:5 | $[M-H]^-$ |
| *824.5442* | PS 39:4 | $[M-H]^-$ |
| *824.5606* | PE O-44:10 / PE P-44:9 | $[M-H]^-$ |
| *825.5657* | PG 40:4 | $[M-H]^-$ |
| *826.4671* | PS 40:10 | $[M-H]^-$ |
| *828.4827* | PS 40:9 | $[M-H]^-$ |
| *829.4879* | PI 34:4 | $[M-H]^-$ |
| *830.4984* | PS 40:8 | $[M-H]^-$ |
| *831.5035* | PI 34:3 | $[M-H]^-$ |
| *832.5140* | PS 40:7 | $[M-H]^-$ |
| *832.5620* | SHexCer 38:2 | $[M-H]^-$ |
| *833.5192* | PI 34:2 | $[M-H]^-$ |
| *834.5297* | PS 40:6 | $[M-H]^-$ |
| *834.5777* | SHexCer 38:1 | $[M-H]^-$ |
| *834.6834* | HexCer 44:3 | $[M-H]^-$ |
| *835.5348* | PI 34:1 | $[M-H]^-$ |
| *836.5242* | PE 44:11 | $[M-H]^-$ |
| *836.5453* | PS 40:5 | $[M-H]^-$ |
| *837.5505* | PI 34:0 | $[M-H]^-$ |
| *838.5398* | PE 44:10 | $[M-H]^-$ |
| *838.5610* | PS 40:4 | $[M-H]^-$ |
| *839.4711* | PI 35:6 | $[M-H]^-$ |
| *840.5555* | PE 44:9 | $[M-H]^-$ |
| *841.4867* | PI 35:5 | $[M-H]^-$ |
| *841.5031* | PG 42:10 | $[M-H]^-$ |
| *843.4436* | DLCL 30:2 | $[M-H]^-$ |
| *848.5569* | SHexCer 38:2 | $[M-H]^-$ |
| *849.5493* | PI 35:1 | $[M-H]^-$ |
| *849.5657* | PG 42:6 | $[M-H]^-$ |
| *850.5726* | SHexCer 38:1 | $[M-H]^-$ |
| *851.4722* | PI 36:7 | $[M-H]^-$ |
| *851.5650* | PI 35:0 | $[M-H]^-$ |
| *851.5814* | PG 42:5 | $[M-H]^-$ |
| *852.4827* | PS 42:11 | $[M-H]^-$ |
| *852.5882* | SHexCer 38:0 | $[M-H]^-$ |

| | | |
|---|---|---|
| *853.4879* | PI 36:6 | $[M-H]^-$ |
| *854.4984* | PS 42:10 | $[M-H]^-$ |
| *855.5035* | PI 36:5 | $[M-H]^-$ |
| *856.5140* | PS 42:9 | $[M-H]^-$ |
| *857.5192* | PI 36:4 | $[M-H]^-$ |
| *858.5297* | PS 42:8 | $[M-H]^-$ |
| *859.5348* | PI 36:3 | $[M-H]^-$ |
| *860.5453* | PS 42:7 | $[M-H]^-$ |
| *860.5933* | SHexCer 40:2 | $[M-H]^-$ |
| *860.6111* | Hex2Cer 34:1 | $[M-H]^-$ |
| *861.5505* | PI 36:2 | $[M-H]^-$ |
| *862.5610* | PS 42:6 | $[M-H]^-$ |
| *862.6090* | SHexCer 40:1 | $[M-H]^-$ |
| *863.5661* | PI 36:1 | $[M-H]^-$ |
| *864.5766* | PS 42:5 | $[M-H]^-$ |
| *864.5882* | SHexCer 39:1 | $[M-H]^-$ |
| *865.4280* | DLCL 32:5 | $[M-H]^-$ |
| *865.5031* | PG 44:12 | $[M-H]^-$ |
| *865.5818* | PI 36:0 | $[M-H]^-$ |
| *867.4436* | DLCL 32:4 | $[M-H]^-$ |
| *867.5024* | PI 37:6 | $[M-H]^-$ |
| *867.5188* | PG 44:11 | $[M-H]^-$ |
| *868.5831* | SHexCer 38:0 | $[M-H]^-$ |
| *869.4593* | DLCL 32:3 | $[M-H]^-$ |
| *869.5180* | PI 37:5 | $[M-H]^-$ |
| *869.5344* | PG 44:10 | $[M-H]^-$ |
| *870.6236* | PS 42:2 | $[M-H]^-$ |
| *871.4749* | DLCL 32:2 | $[M-H]^-$ |
| *871.5337* | PI 37:4 | $[M-H]^-$ |
| *873.5493* | PI 37:3 | $[M-H]^-$ |
| *873.5657* | PG 44:8 | $[M-H]^-$ |
| *874.6090* | SHexCer 41:2 | $[M-H]^-$ |
| *876.5882* | SHexCer 40:2 | $[M-H]^-$ |
| *876.6246* | SHexCer 41:1 | $[M-H]^-$ |
| *877.5806* | PI 37:1 | $[M-H]^-$ |
| *877.5970* | PG 44:6 | $[M-H]^-$ |
| *878.4984* | PS 44:12 | $[M-H]^-$ |
| *878.6039* | SHexCer 40:1 | $[M-H]^-$ |
| *879.5035* | PI 38:7 | $[M-H]^-$ |
| *879.5963* | PI 37:0 | $[M-H]^-$ |
| *879.6127* | PG 44:5 | $[M-H]^-$ |
| *880.5140* | PS 44:11 | $[M-H]^-$ |

| | | |
|---|---|---|
| *880.6195* | SHexCer 40:0 | [M-H]- |
| *881.5192* | PI 38:6 | [M-H]- |
| *882.5297* | PS 44:10 | [M-H]- |
| *883.5348* | PI 38:5 | [M-H]- |
| *884.5453* | PS 44:9 | [M-H]- |
| *885.5505* | PI 38:4 | [M-H]- |
| *886.5610* | PS 44:8 | [M-H]- |
| *886.6090* | SHexCer 42:3 | [M-H]- |
| *887.5661* | PI 38:3 | [M-H]- |
| *888.5766* | PS 44:7 | [M-H]- |
| *888.6246* | SHexCer 42:2 | [M-H]- |
| *889.5818* | PI 38:2 | [M-H]- |
| *890.6403* | SHexCer 42:1 | [M-H]- |
| *891.4436* | DLCL 34:6 | [M-H]- |
| *892.6079* | PS 44:5 | [M-H]- |
| *892.6195* | SHexCer 41:1 | [M-H]- |
| *893.4593* | DLCL 34:5 | [M-H]- |
| *893.6131* | PI 38:0 | [M-H]- |
| *894.6352* | SHexCer 41:0 | [M-H]- |
| *895.5337* | PI 39:6 | [M-H]- |
| *897.5493* | PI 39:5 | [M-H]- |
| *899.5062* | DLCL 34:2 | [M-H]- |
| *899.5650* | PI 39:4 | [M-H]- |
| *900.6246* | SHexCer 43:3 | [M-H]- |
| *902.6039* | SHexCer 42:3 | [M-H]- |
| *902.6403* | SHexCer 43:2 | [M-H]- |
| *903.5963* | PI 39:2 | [M-H]- |
| *904.6195* | SHexCer 42:2 | [M-H]- |
| *905.5192* | PI 40:8 | [M-H]- |
| *905.6119* | PI 39:1 | [M-H]- |
| *906.6352* | SHexCer 42:1 | [M-H]- |
| *907.5348* | PI 40:7 | [M-H]- |
| *907.6276* | PI 39:0 | [M-H]- |
| *908.6508* | SHexCer 42:0 | [M-H]- |
| *909.5505* | PI 40:6 | [M-H]- |
| *911.5661* | PI 40:5 | [M-H]- |
| *912.5366* | SHex2Cer 32:1 | [M-H]- |
| *913.5818* | PI 40:4 | [M-H]- |
| *915.4436* | DLCL 36:8 | [M-H]- |
| *916.6559* | SHexCer 44:2 | [M-H]- |
| *916.6737* | Hex2Cer 38:1 | [M-H]- |
| *917.4593* | DLCL 36:7 | [M-H]- |

| | | |
|---|---|---|
| *918.6352* | SHexCer 43:2 | $[M-H]^-$ |
| *919.4749* | DLCL 36:6 | $[M-H]^-$ |
| *919.6287* | PI 40:1 | $[M-H]^-$ |
| *920.6508* | SHexCer 43:1 | $[M-H]^-$ |
| *921.6444* | PI 40:0 | $[M-H]^-$ |
| *925.5219* | DLCL 36:3 | $[M-H]^-$ |
| *927.5035* | PI 42:11 | $[M-H]^-$ |
| *929.5192* | PI 42:10 | $[M-H]^-$ |
| *932.6508* | SHexCer 44:2 | $[M-H]^-$ |
| *934.6665* | SHexCer 44:1 | $[M-H]^-$ |
| *938.5522* | SHex2Cer 34:2 | $[M-H]^-$ |
| *940.5679* | SHex2Cer 34:1 | $[M-H]^-$ |
| *941.4593* | DLCL 38:9 | $[M-H]^-$ |
| *945.4906* | DLCL 38:7 | $[M-H]^-$ |
| *947.5062* | DLCL 38:6 | $[M-H]^-$ |
| *949.5219* | DLCL 38:5 | $[M-H]^-$ |
| *957.5845* | DLCL 38:1 | $[M-H]^-$ |
| *959.6001* | DLCL 38:0 | $[M-H]^-$ |
| *968.5992* | SHex2Cer 36:1 | $[M-H]^-$ |
| *969.4906* | DLCL 40:9 | $[M-H]^-$ |
| *993.4906* | DLCL 42:11 | $[M-H]^-$ |
| *1019.5062* | DLCL 44:12 | $[M-H]^-$ |
| *1021.5219* | DLCL 44:11 | $[M-H]^-$ |
| *1179.7372* | NeuAcHex2Cer 36:1 | $[M-H]^-$ |
| *1207.7685* | NeuAcHex2Cer 38:1 | $[M-H]^-$ |
| *1382.8166* | Hex(2)-HexNAc-NeuAc-Cer 36:1 | $[M-H]^-$ |
| *1410.8479* | Hex(2)-HexNAc-NeuAc-Cer 38:1 | $[M-H]^-$ |
| *1470.8326* | NeuAc2Hex2Cer 36:1 | $[M-H]^-$ |
| *1544.8694* | Hex(3)-HexNAc-NeuAc-Cer 36:1 | $[M-H]^-$ |
| *1572.9007* | Hex(3)-HexNAc-NeuAc-Cer 38:1 | $[M-H]^-$ |
| *1600.9320* | Hex(3)-HexNAc-NeuAc-Cer 40:1 | $[M-H]^-$ |
| *1690.9273* | Hex(3)-HexNAc-Fuc-NeuAc-Cer 36:1 | $[M-H]^-$ |
| *1734.9535* | Hex(3)-HexNAc-Fuc-NeuGc-Cer 38:1/Hex(4)-HexNAc-NeuAc-Cer 38:1 | $[M-H]^-$ |
| *1747.9488* | Hex(3)-HexNAc(2)-NeuAc-Cer 36:1 | $[M-H]^-$ |
| *1775.9801* | Hex(3)-HexNAc(2)-NeuAc-Cer 38:1 | $[M-H]^-$ |
| *1781.9430* | Hex(3)-HexNAc-KDN(2)-Cer 38:1 | $[M-H]^-$ |
| *1789.9593* | Hex(2)-NeuAc(3)-Cer 38:1 | $[M-H]^-$ |
| *1791.9386* | Hex(2)-HexNAc-Fuc-NeuAc(2)-Cer 34:1 | $[M-H]^-$ |
| *1791.9750* | Hex(3)-HexNAc(2)-NeuGc-Cer 38:1 | $[M-H]^-$ |
| *1817.9906* | Hex(2)-NeuAc(3)-Cer 40:1 | $[M-H]^-$ |
| *1819.9699* | Hex(2)-HexNAc-Fuc-NeuAc(2)-Cer 36:1 | $[M-H]^-$ |

| | | |
|---|---|---|
| *1820.0063* | Hex(3)-HexNAc(2)-NeuGc-Cer 40:1 | [M-H]$^-$ |
| *1835.9648* | Hex(3)-HexNAc-NeuAc(2)-Cer 36:1 | [M-H]$^-$ |
| *1835.9899* | Hex(3)-HexNAc-KDN(2)-Cer 42:2 | [M-H]$^-$ |
| *1838.0056* | Hex(3)-HexNAc-KDN(2)-Cer 42:1 | [M-H]$^-$ |
| *1863.9961* | Hex(3)-HexNAc-NeuAc(2)-Cer 38:1 | [M-H]$^-$ |
| *1864.0212* | Hex(3)-HexNAc-KDN(2)-Cer 44:2 | [M-H]$^-$ |
| *1865.9754* | Hex(3)-HexNAc(2)-Fuc-NeuAc-Cer 34:1 | [M-H]$^-$ |
| *1866.0369* | Hex(3)-HexNAc-KDN(2)-Cer 44:1 | [M-H]$^-$ |
| *1879.991* | Hex(3)-HexNAc-NeuAc-NeuGc-Cer 38:1 | [M-H]$^-$ |
| *2083.0704* | Hex(3)-HexNAc(2)-NeuAc-NeuGc-Cer 38:1 | [M-H]$^-$ |
| *2084.0908* | Hex(4)-HexNAc(2)-Fuc-NeuAc-Cer 38:1 | [M-H]$^-$ |
| *2127.0602* | Hex(3)-HexNAc-NeuAc(3)-Cer 36:1 | [M-H]$^-$ |
| *2129.0759* | Hex(4)-HexNAc(3)-NeuGc-Cer 36:1 | [M-H]$^-$ |
| *2155.0915* | Hex(3)-HexNAc-NeuAc(3)-Cer 38:1 | [M-H]$^-$ |

**Supplementary Table 2. In-house curated lipid database of commonly observed negative lipid ions and expansive theoretical mass ganglioside panel.**

| | Genotype | | |
|---|---|---|---|
| | *Sgsh*$^{+/+}$ ♀ | *Sgsh*$^{+/R245H}$ ♀ | *Sgsh*$^{R245H/R245H}$ ♀ |
| | *Sgsh*$^{+/+}$ ♂ | *Sgsh*$^{+/R245H}$ ♂ | *Sgsh*$^{R245H/R245H}$ ♂ |
| **Median pair-to-birth interval (days)** | 20 (range 19-22) | 24 (range 19-42) | 21 (range 20-25) |
| **Median litter size at birth** | 8 (range 6-9) | 7.5 (range 1-11) | 7 (range 5-8) |
| **Median litter size at weaning** | 8 (range 6-9) | 6.5 (range 1-11) | 6 (range 5-8) |
| **Entire litters lost before counting (% total number of pairs)** | 9% (1/11 pairs) | 0% (0/30 pairs) | 11% (2/19 pairs) |

**Supplementary Table 3. Breeding statistics from combinations of genotype pairs.**

| *m/z* | **Putative Identity** | **Genotype (F)** | **Genotype (adj. p)** | **Colony (F)** | **Colony (adj. p)** | **Interaction (F)** | **Interaction (adj. p)** |
|---|---|---|---|---|---|---|---|
| 806.5464 | SHexCer 36:1;O2 | 18.04 | 0.0353 | 0.62 | 0.75 | 0.0420 | 0.99 |
| 820.5256 | SHexCer 36:2;O3 | 18.39 | 0.0353 | 0.15 | 0.83 | 0.19 | 0.99 |
| 822.5413 | SHexCer 36:1;O3 | 30.86 | 0.0135 | 0.60 | 0.75 | 0.0320 | 0.99 |
| 865.5031 | BMP 44:12 | 33.93 | 0.0133 | 1.97 | 0.75 | 0.0500 | 0.99 |
| 867.5188 | BMP 44:11 | 21.78 | 0.0267 | 1.45 | 0.75 | 0.0690 | 0.99 |
| 968.5992 | SHex2Cer 36:1;O2 | 133.98 | 0.0003 | 0.26 | 0.79 | 0.19 | 0.99 |
| 1179.7372 | $G_{M3}$ 36:1 | 76.80 | 0.0011 | 0.22 | 0.79 | 0.0120 | 0.99 |
| 1207.7685 | $G_{M3}$ 38:1 | 25.93 | 0.0167 | 0.0090 | 0.95 | 0.0010 | 0.99 |
| 1382.8166 | $G_{M2}$ 36:1 | 231.59 | 0.0001 | 0.13 | 0.83 | 0.0040 | 0.99 |
| 1410.8479 | $G_{M2}$ 38:1 | 14.72 | 0.0500 | 0.33 | 0.79 | 0.20 | 0.99 |
| 1470.8326 | $G_{D3}$ 36:1 | 100.80 | 0.0005 | 0.31 | 0.79 | 0.0009 | 0.99 |

**Supplementary Table 4. Lipid species significantly elevated in MPS IIIA mice, determined by univariate two-way ANOVA.** F-values (F) and adjusted p-values (adj. p) are shown for genotype, colony, and interaction effects. Statistically significant differences determined as adj. p≤0.05. BMP, bis(monoacylglycero)phosphate; $G_{M3}$, monosialodihexosylganglioside; $G_{M2}$, monosialotrihexosylganglioside; $G_{D3}$, disialodihexosylganglioside.

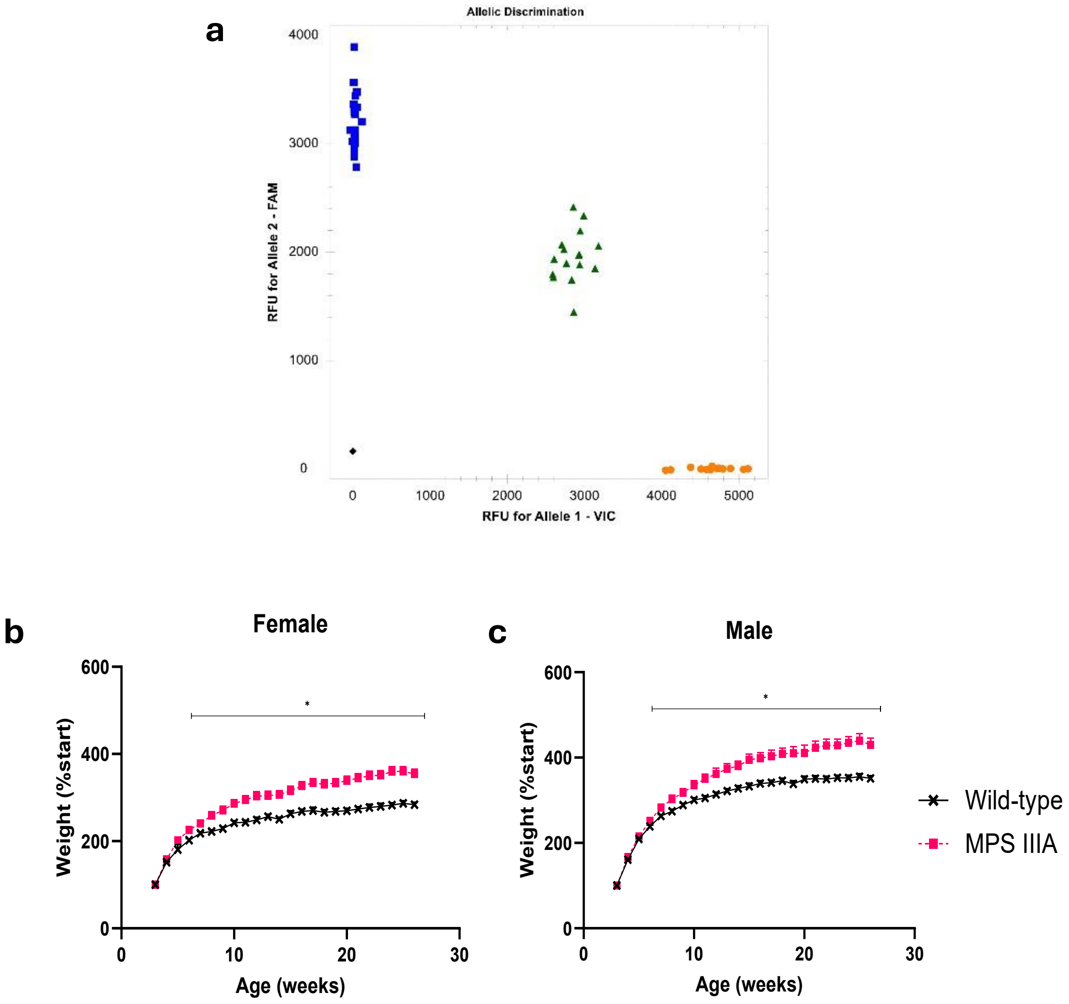


**Supplementary Figure 1: Generation of the *Sgsh*$^{R245H}$ variant in mice by germline CRISPR/Cas9 gene targeting.** (a) Example of an allelic discrimination plot used to assign genotypes showing wild-type (VIC-label, orange circles), MPS IIIA (FAM-label, blue squares) or heterozygote status (VIC- and FAM-label, green triangles) of mice. A control without template DNA is also included (black diamond). (b, c) The percentage change in mean body weight in wild-type and MPS IIIA female and male mice is shown (n=16 mice/group). Statistically significant differences between sex-matched genotype groups were found from 6 weeks of age onwards. *$p \leq 0.05$.

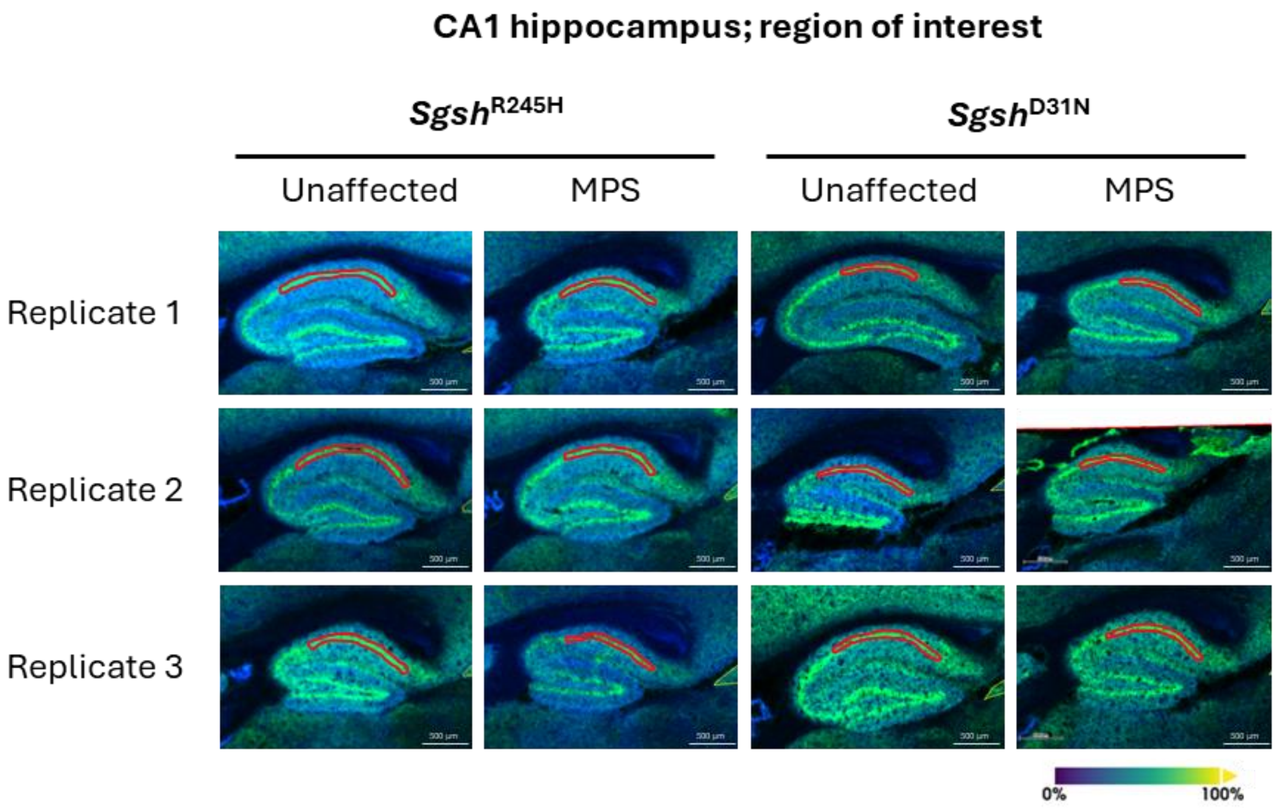


**Supplementary Figure 2: Mass spectrometry images of the CA1 hippocampal region of interest in MPS IIIA and unaffected mice.** Brain sections reflect MPS IIIA mice expressing the *Sgsh*$^{R245H}$ (CRISPR/Cas9 gene targeting) or *Sgsh*$^{D31N}$ (spontaneous) variants, or strain-matched unaffected controls (n=3/group) at 20 weeks of age. CA1 regions of interest were defined by selecting signals corresponding to neutral (unchanged) putative phospholipids, phosphatidylethanolamine (PE) 38:4 (*m/z* 766.5398) and phosphatidylinositol (PI) 38:5 (*m/z* 883.5348), followed by manual delineation on each section, indicated by a red outline.

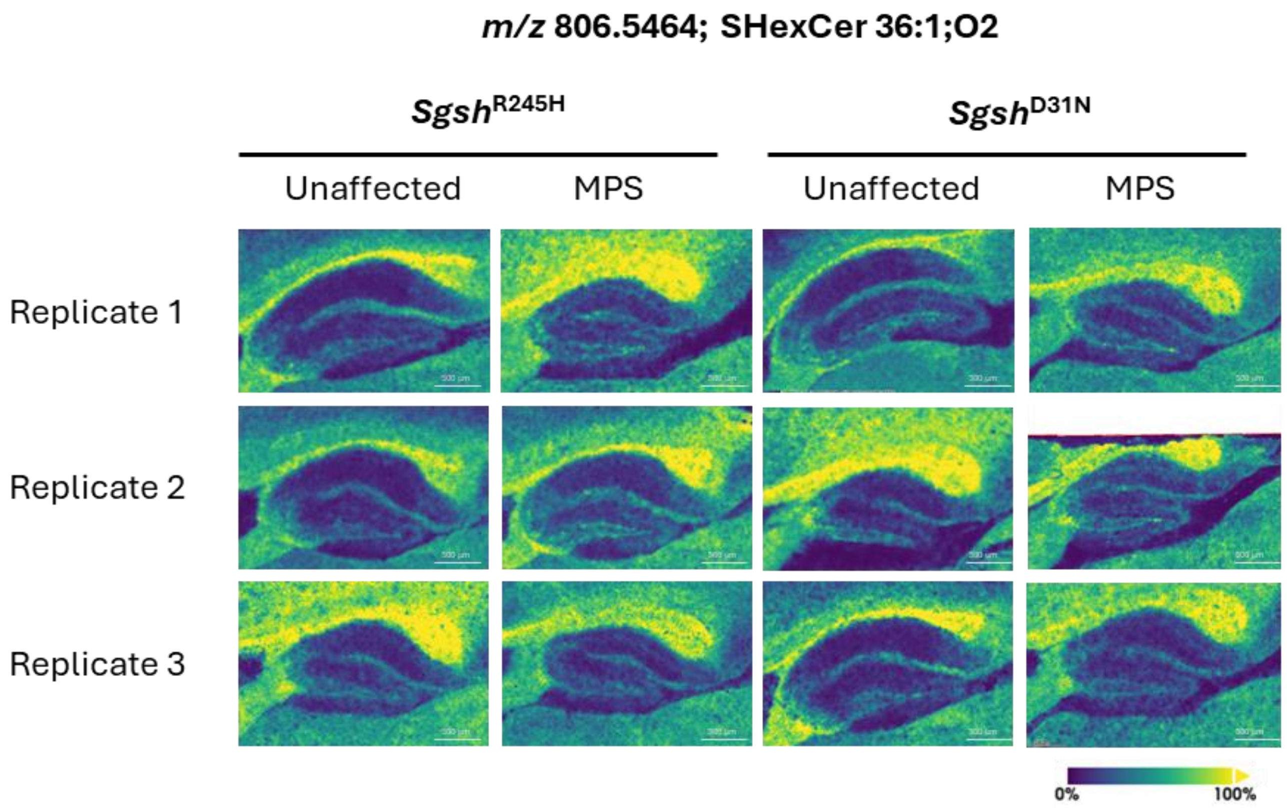


**Supplementary Figure 3: Mass spectrometry images depicting *m/z* 806.5464 signal intensity in MPS IIIA and unaffected mice.** Brain sections reflect MPS IIIA mice expressing the *Sgsh*$^{R245H}$ (CRISPR/Cas9 gene targeting) or *Sgsh*$^{D31N}$ (spontaneous) variants, or strain-matched unaffected controls (n=3/group) at 20 weeks of age. *m/z* 806.5464 was putatively assigned as SHexCer 36:1;O2.

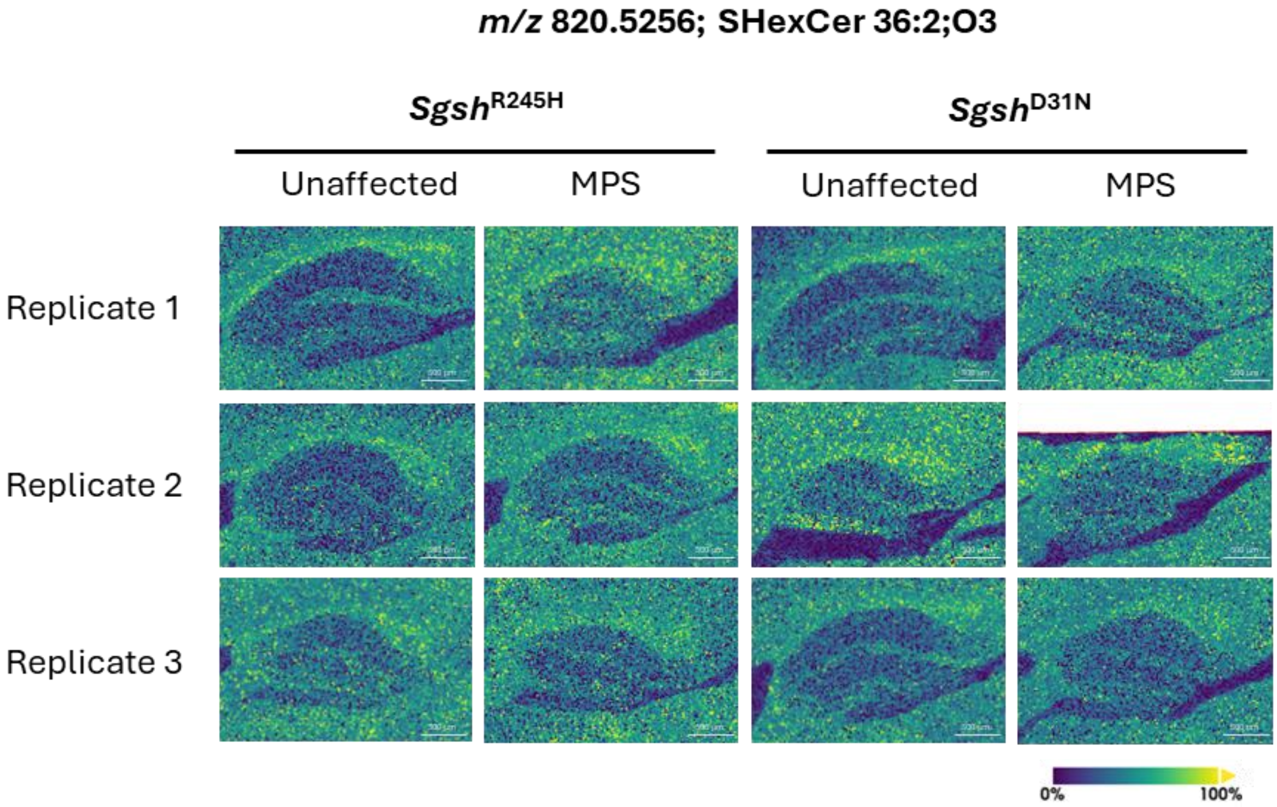


**Supplementary Figure 4: Mass spectrometry images depicting *m/z* 820.5256 signal intensity in MPS IIIA and unaffected mice.** Brain sections reflect MPS IIIA mice expressing the $\textit{Sgsh}^{R245H}$ (CRISPR/Cas9 gene targeting) or $\textit{Sgsh}^{D31N}$ (spontaneous) variants, or strain-matched unaffected controls (n=3/group) at 20 weeks of age. *m/z* 820.5256 was putatively assigned as SHexCer 36:2;O3.

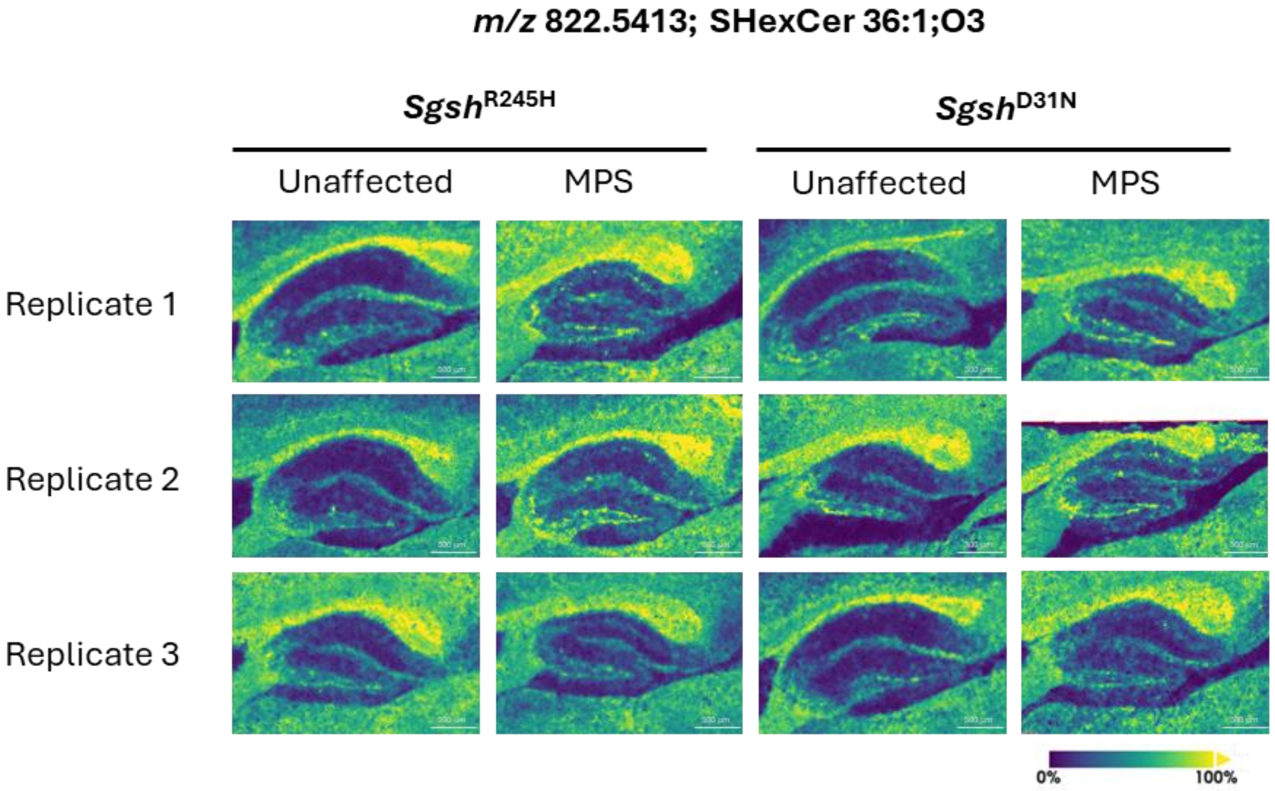


**Supplementary Figure 5: Mass spectrometry images depicting *m/z* 822.5413 signal intensity in MPS IIIA and unaffected mice.** Brain sections reflect MPS IIIA mice expressing the *Sgsh*$^{R245H}$ (CRISPR/Cas9 gene targeting) or *Sgsh*$^{D31N}$ (spontaneous) variants, or strain-matched unaffected controls (n=3/group) at 20 weeks of age. *m/z* 822.5413 was putatively assigned as SHexCer 36:1;O3.

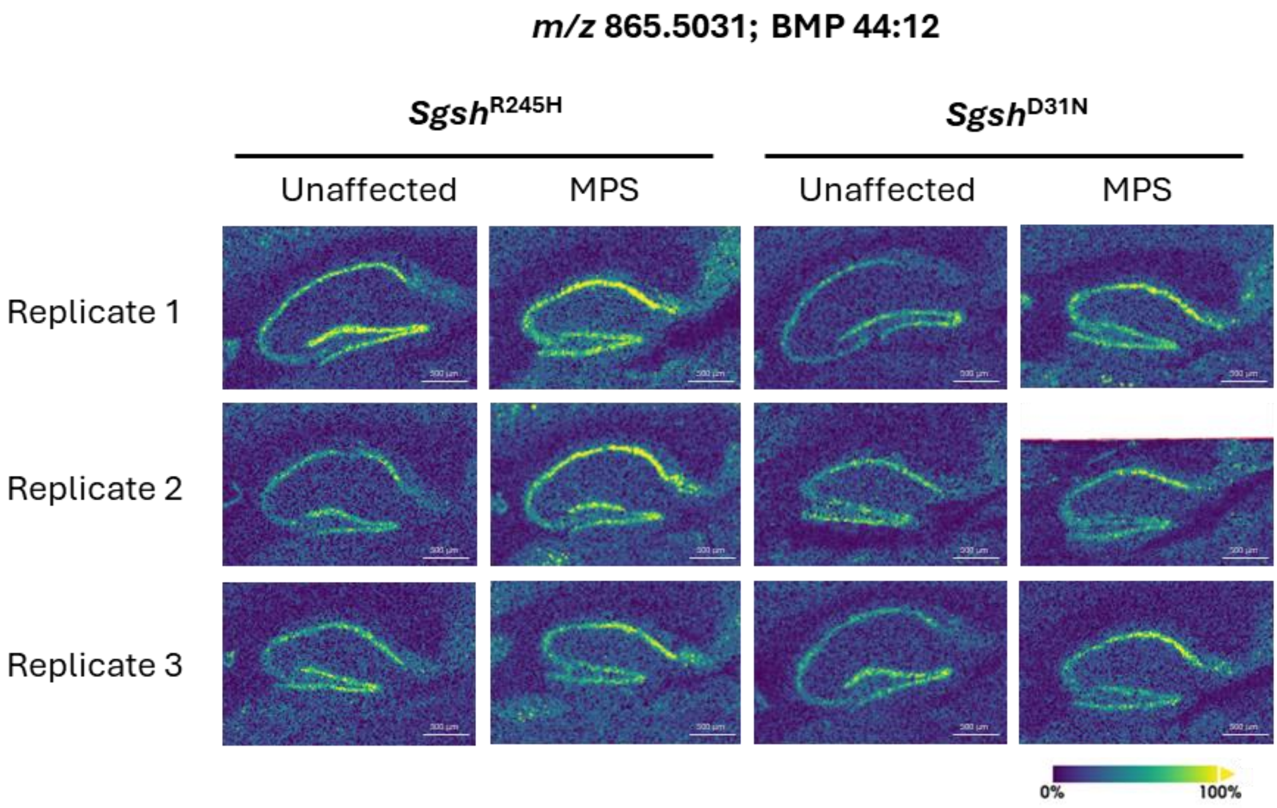


**Supplementary Figure 6: Mass spectrometry images depicting *m/z* 865.5031 signal intensity in MPS IIIA and unaffected mice.** Brain sections reflect MPS IIIA mice expressing the *Sgsh*$^{R245H}$ (CRISPR/Cas9 gene targeting) or *Sgsh*$^{D31N}$ (spontaneous) variants, or strain-matched unaffected controls (n=3/group) at 20 weeks of age. *m/z* 865.5031 was putatively assigned as bis(monoacylglycero)phosphate (BMP) 44:12.

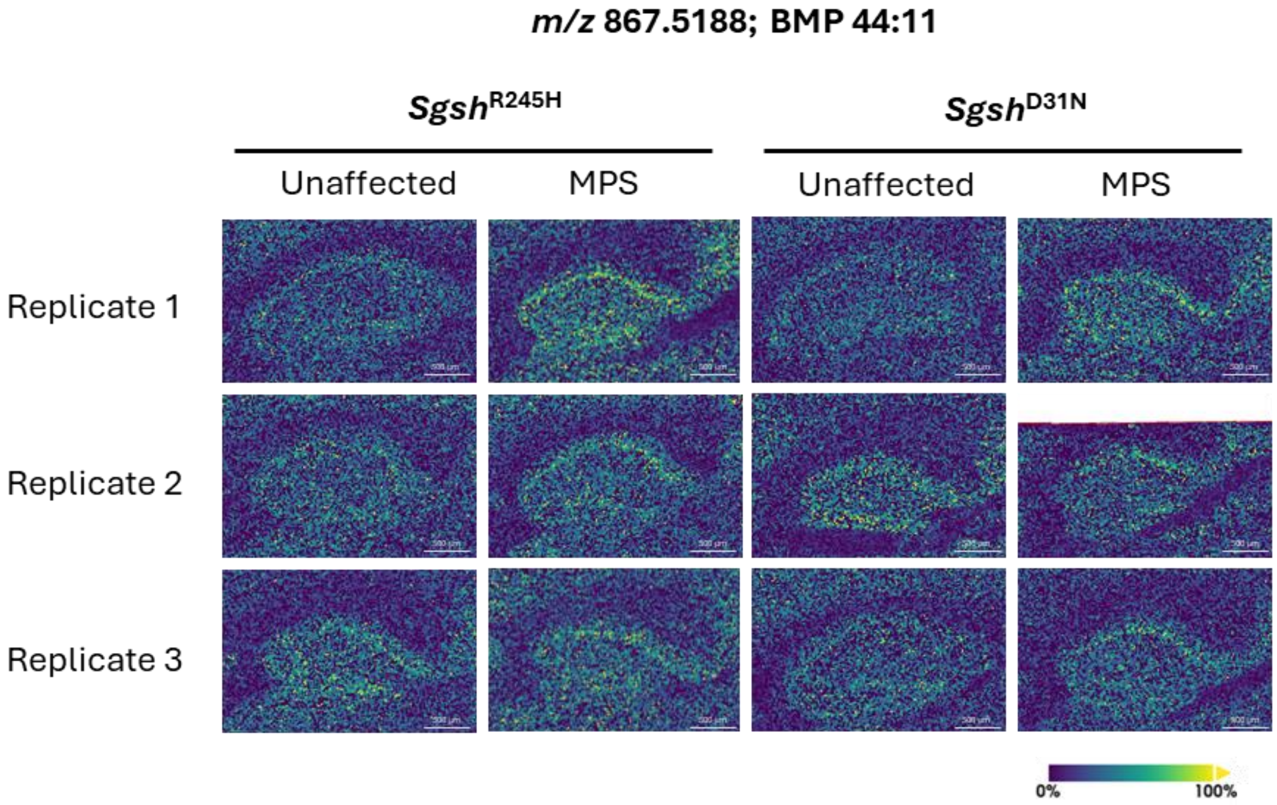


**Supplementary Figure 7: Mass spectrometry images depicting *m/z* 867.5188 signal intensity in MPS IIIA and unaffected mice.** Brain sections reflect MPS IIIA mice expressing the *Sgsh*$^{R245H}$ (CRISPR/Cas9 gene targeting) or *Sgsh*$^{D31N}$ (spontaneous) variants, or strain-matched unaffected controls (n=3/group) at 20 weeks of age. *m/z* 867.5188 was putatively assigned as bis(monoacylglycero)phosphate (BMP) 44:11.

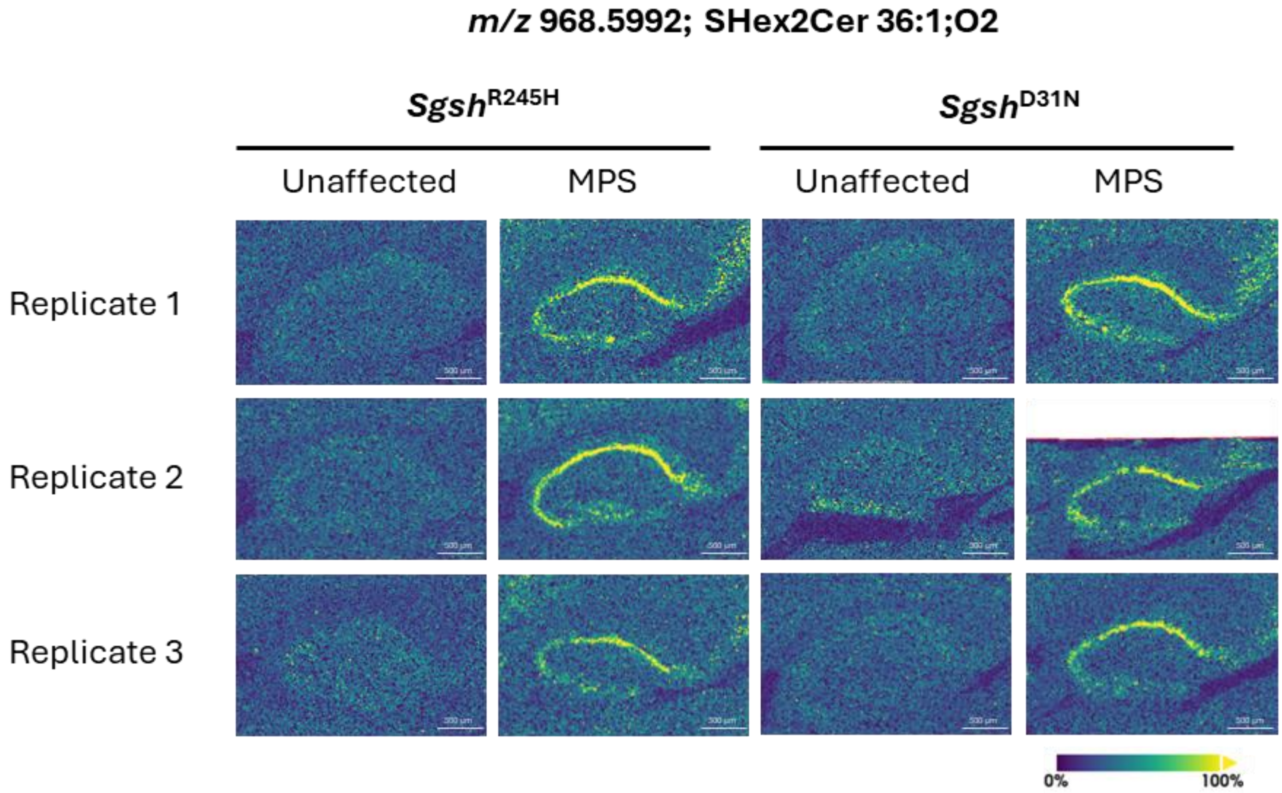


**Supplementary Figure 8: Mass spectrometry images depicting *m/z* 968.5992 signal intensity in MPS IIIA and unaffected mice.** Brain sections reflect MPS IIIA mice expressing the *Sgsh*$^{R245H}$ (CRISPR/Cas9 gene targeting) or *Sgsh*$^{D31N}$ (spontaneous) variants, or strain-matched unaffected controls (n=3/group) at 20 weeks of age. *m/z* 968.5992 was putatively assigned as SHex2Cer 36:1;O2.

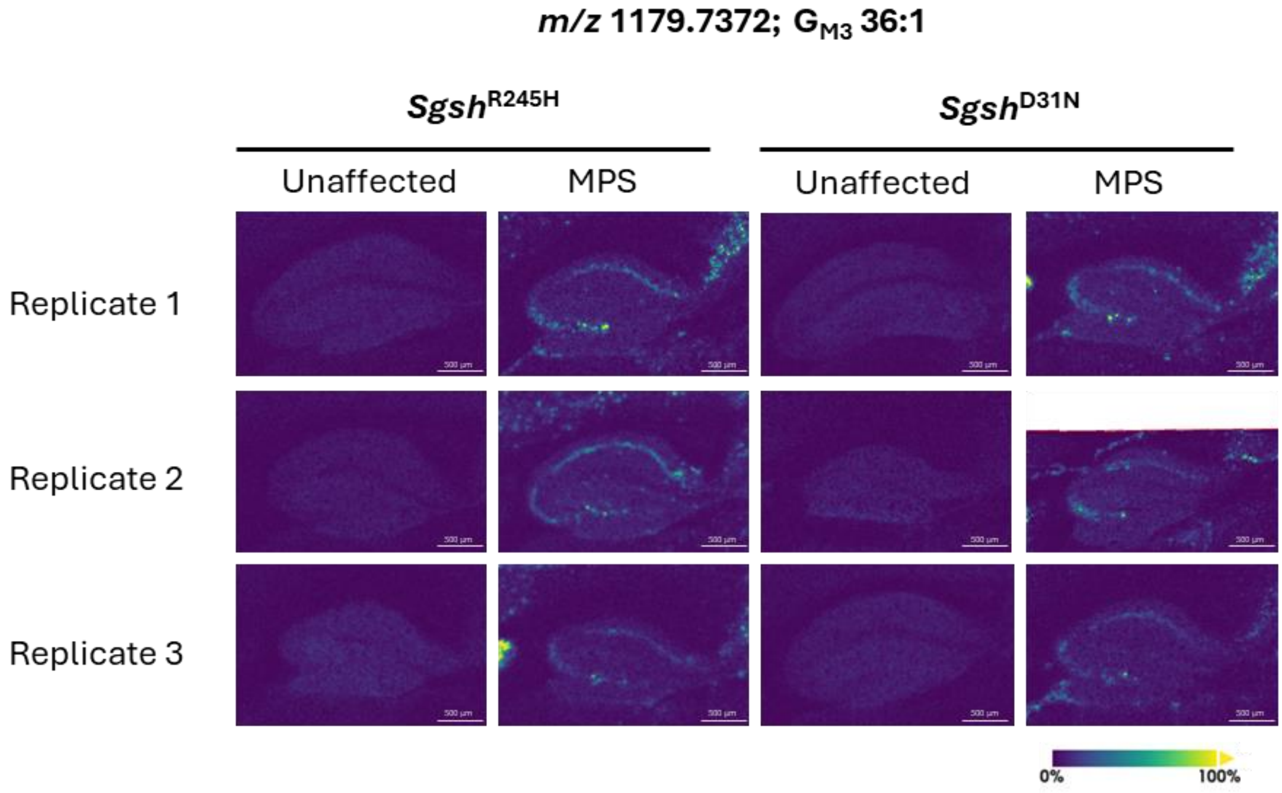


**Supplementary Figure 9: Mass spectrometry images depicting *m/z* 1179.7372 signal intensity in MPS IIIA and unaffected mice.** Brain sections reflect MPS IIIA mice expressing the *Sgsh*$^{R245H}$ (CRISPR/Cas9 gene targeting) or *Sgsh*$^{D31N}$ (spontaneous) variants, or strain-matched unaffected controls (n=3/group) at 20 weeks of age. *m/z* 1179.7372 was putatively assigned as monosialodihexosylganglioside ($G_{M3}$) 36:1.

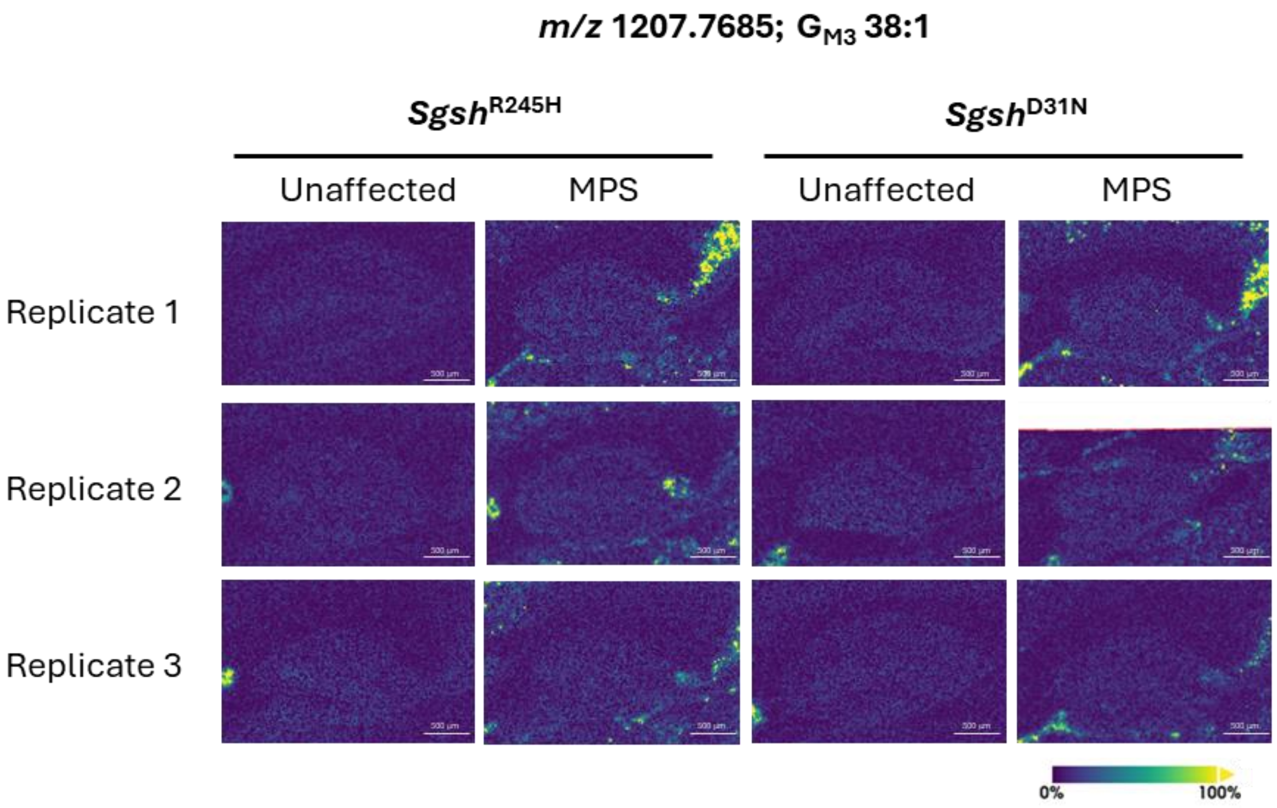


**Supplementary Figure 10: Mass spectrometry images depicting *m/z* 1207.7685 signal intensity in MPS IIIA and unaffected mice.** Brain sections reflect MPS IIIA mice expressing the *Sgsh*$^{R245H}$ (CRISPR/Cas9 gene targeting) or *Sgsh*$^{D31N}$ (spontaneous) variants, or strain-matched unaffected controls (n=3/group) at 20 weeks of age. *m/z* 1207.7685 was putatively assigned as monosialodihexosylganglioside ($G_{M3}$) 38:1.

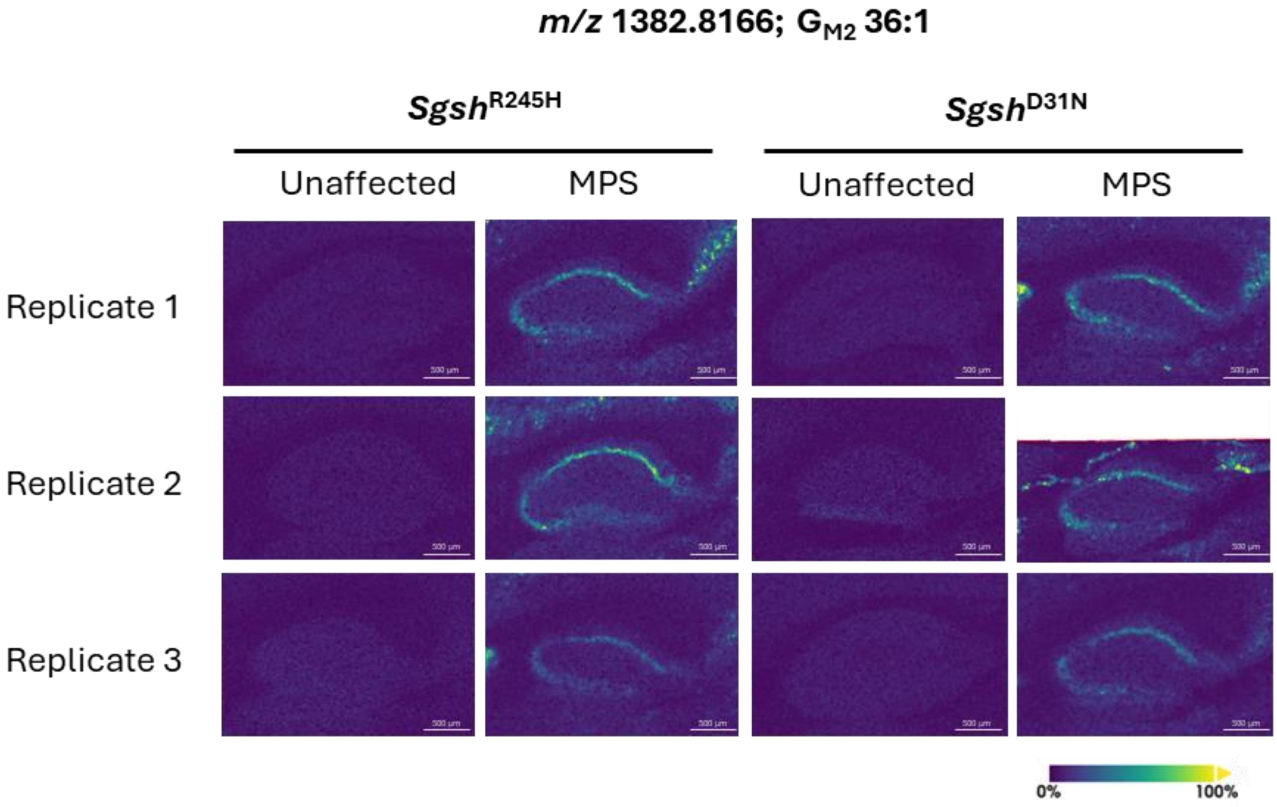


**Supplementary Figure 11: Mass spectrometry images depicting *m/z* 1382.8166 signal intensity in MPS IIIA and unaffected mice.** Brain sections reflect MPS IIIA mice expressing the *Sgsh*$^{R245H}$ (CRISPR/Cas9 gene targeting) or *Sgsh*$^{D31N}$ (spontaneous) variants, or strain-matched unaffected controls (n=3/group) at 20 weeks of age. *m/z* 1382.8166 was putatively assigned as monosialotrihexosylganglioside ($G_{M2}$) 36:1.

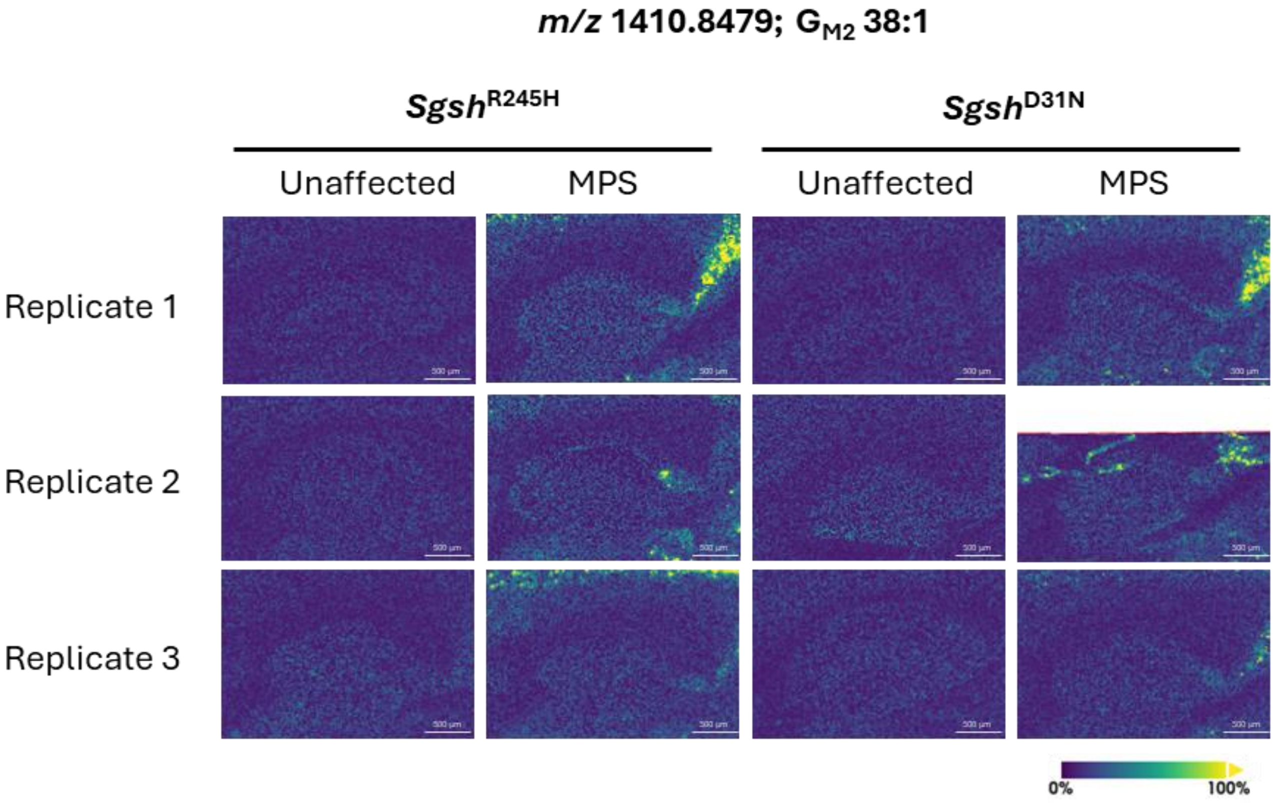


**Supplementary Figure 12: Mass spectrometry images depicting *m/z* 1410.8479 signal intensity in MPS IIIA and unaffected mice.** Brain sections reflect MPS IIIA mice expressing the $Sgsh^{R245H}$ (CRISPR/Cas9 gene targeting) or $Sgsh^{D31N}$ (spontaneous) variants, or strain-matched unaffected controls (n=3/group) at 20 weeks of age. *m/z* 1410.8479 was putatively assigned as monosialotrihexosylganglioside ($G_{M2}$) 38:1.

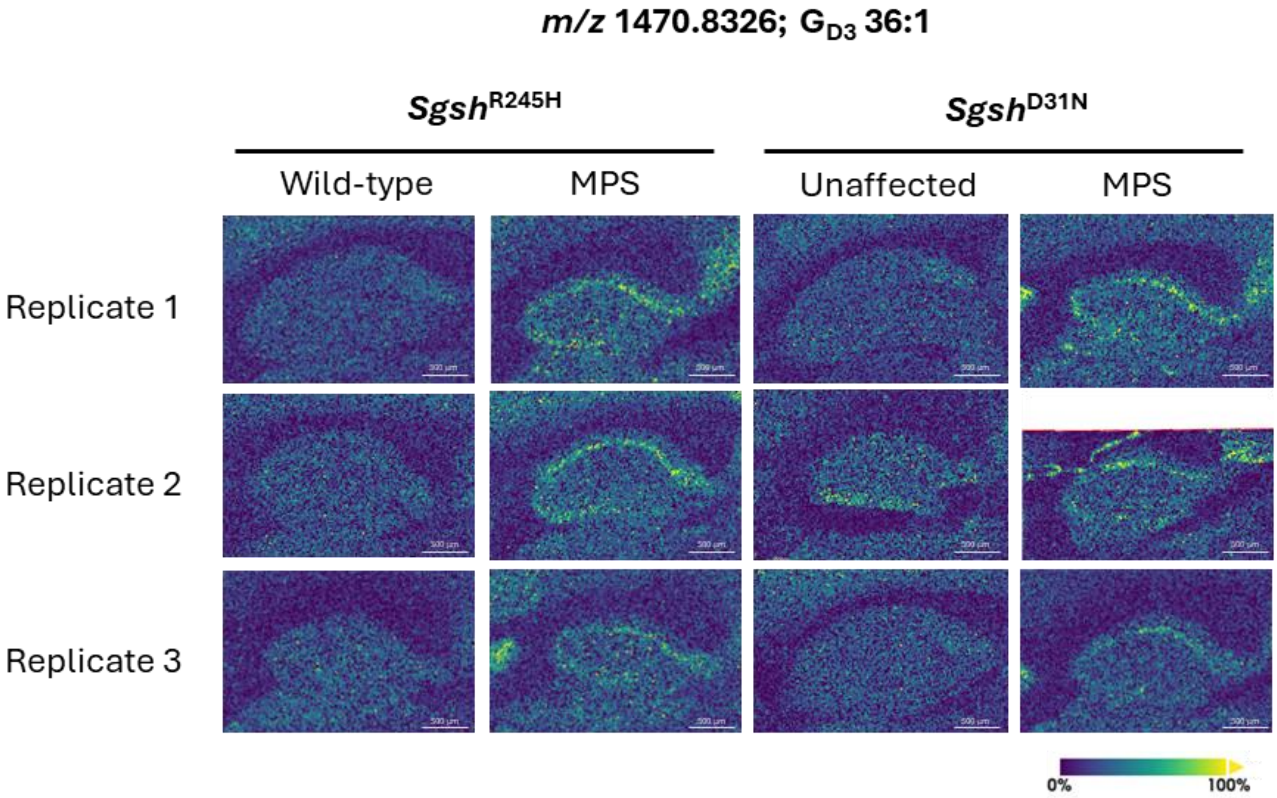


**Supplementary Figure 13: Mass spectrometry images depicting *m/z* 1470.8326 signal intensity in MPS IIIA and unaffected mice.** Brain sections reflect MPS IIIA mice expressing the *Sgsh*$^{R245H}$ (CRISPR/Cas9 gene targeting) or *Sgsh*$^{D31N}$ (spontaneous) variants, or strain-matched unaffected controls (n=3/group) at 20 weeks of age. *m/z* 1470.8326 was putatively assigned as disialodihexosylganglioside ($G_{D3}$) 36:1.

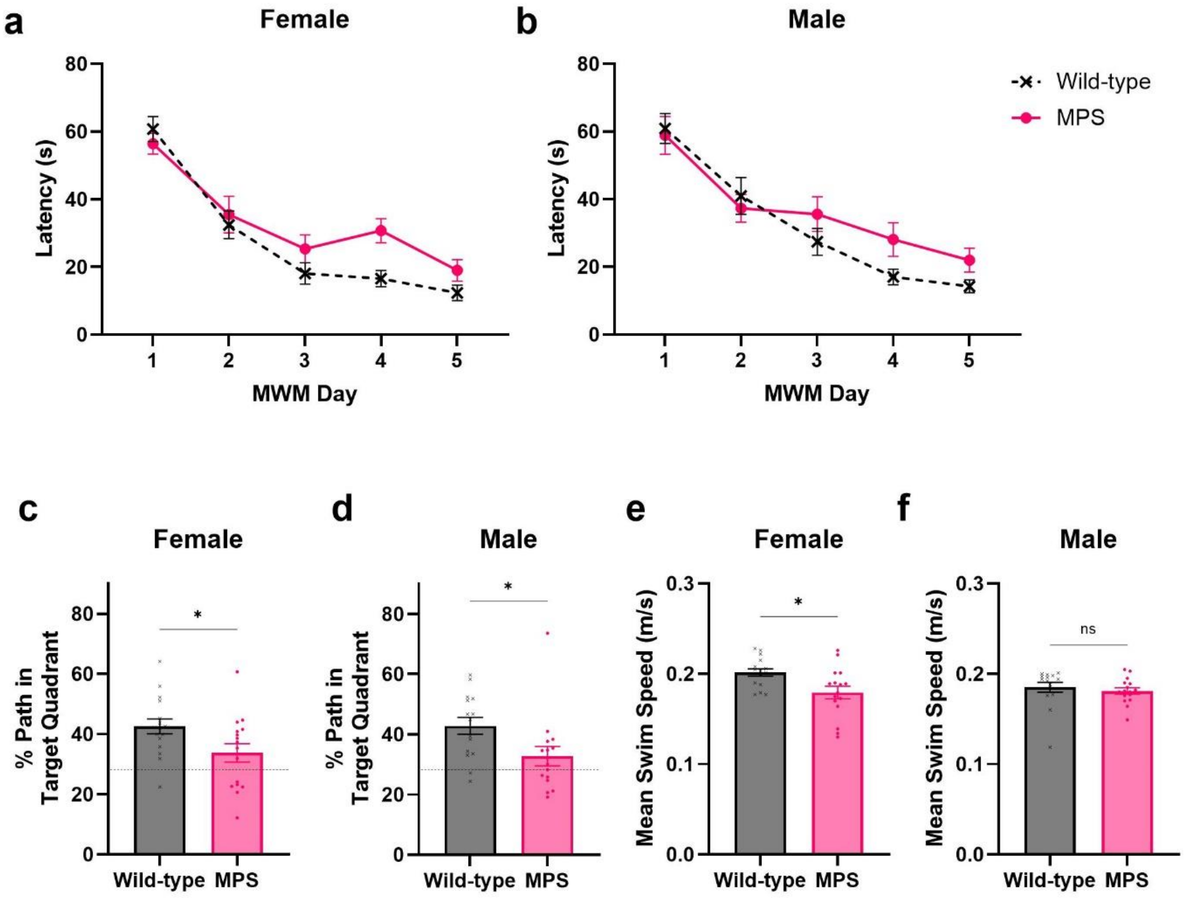


**Supplementary Figure 14: MPS IIIA mice exhibit memory and learning deficits in the Morris Water Maze probe phase.** (a,b) The average swim time to find the hidden platform in the Acquisition Phase of the Morris Water Maze is shown for 22-23-week-old female and male mice. (c-f) For the Probe Phase (day 6 of testing), the platform was removed from the pool, and the exploration pattern was recorded. The dashed line indicates the 25% chance level if no preference is shown. The percentage of (c,d) path length spent in the target quadrant was analysed with ANY-Maze software. (e.f) Average swim speeds of mice in the 90-second probe test. n=15-16 mice/group; the data from one male wild-type mouse was excluded due to failing the visual test on day 7. ns, not significant; *$p \leq 0.05$.

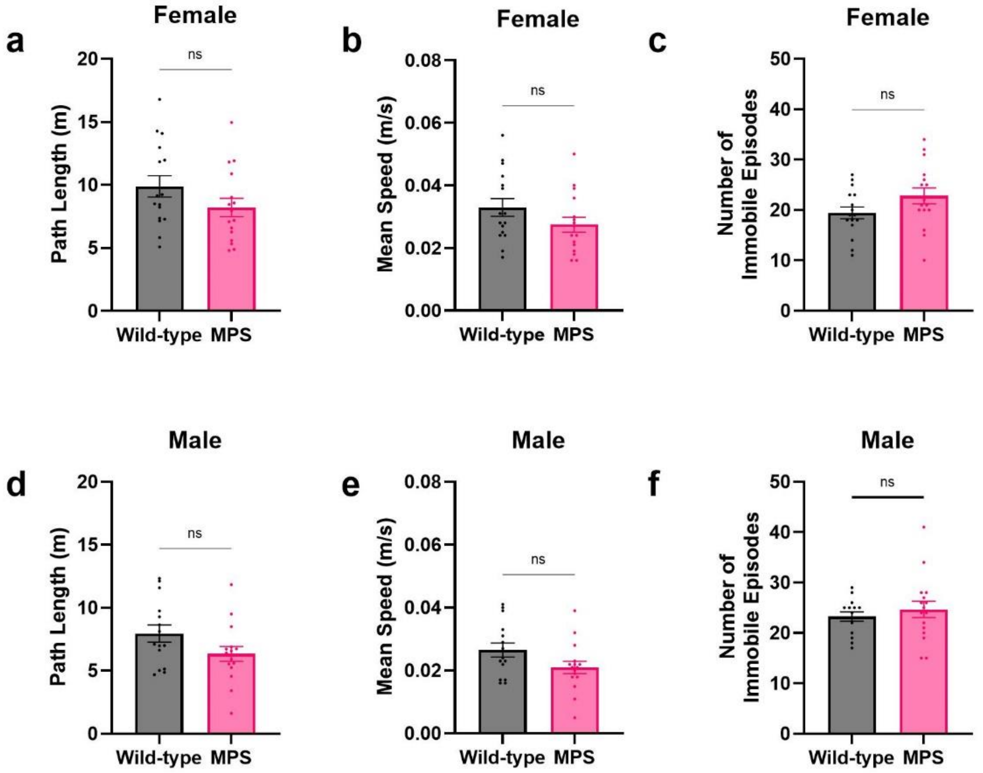


**Supplementary Figure 15: Locomotor activity is not impacted by genotype in 23-week-old mice evaluated in the open field as part of a behavioural test battery.** Wild-type and MPS IIIA (a,b,c) female and (d,e,f) male mice were tested in an open field arena with dimensions of 30 x 50 cm. The exploration over 5 minutes was analysed using ANY-Maze software. The parameters that were assessed included (a,d) path length travelled; (b,e) average speed during exploration and (c,f) the number of immobile freezing episodes. n=16 mice/group. ns, not significant.

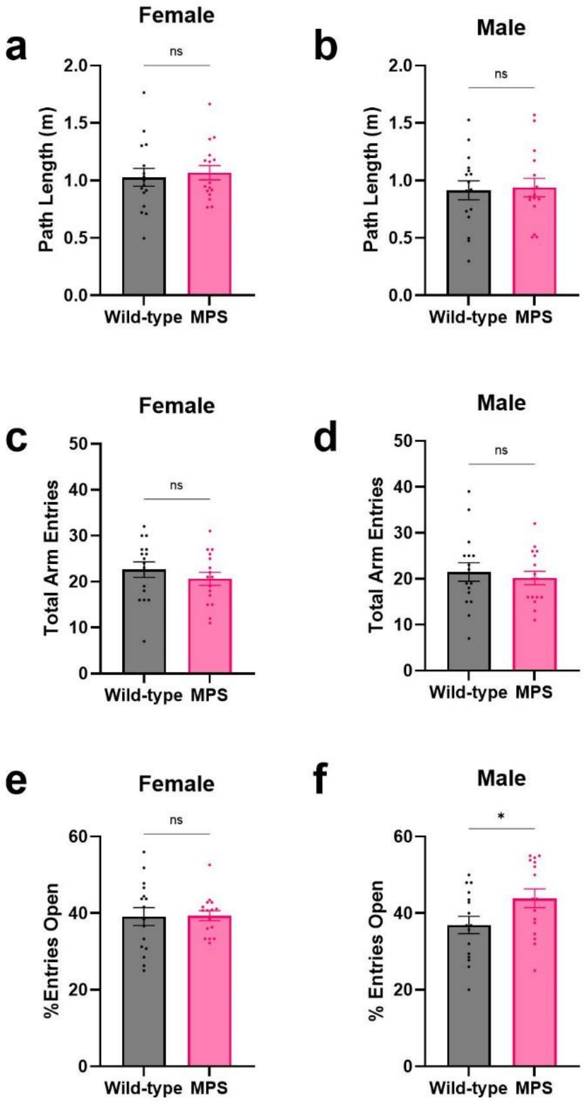


**Supplementary Figure 16: Evaluation of anxiety-related behaviours in the Elevated Plus Maze.** The exploration over 5 minutes in the Elevated Plus Maze was recorded in wild-type and MPS IIIA mice at 24 weeks of age. Female (a, c, e) and male (b, d, f) mice were both evaluated. The assessed parameters included the (a,b) total path length; (c,d) number of entries into open and closed arms (total arm entries); (e,f) the number of open arm entries as a percentage of total arm entries. n=16 mice/group. ns, not significant; *$p \leq 0.05$.

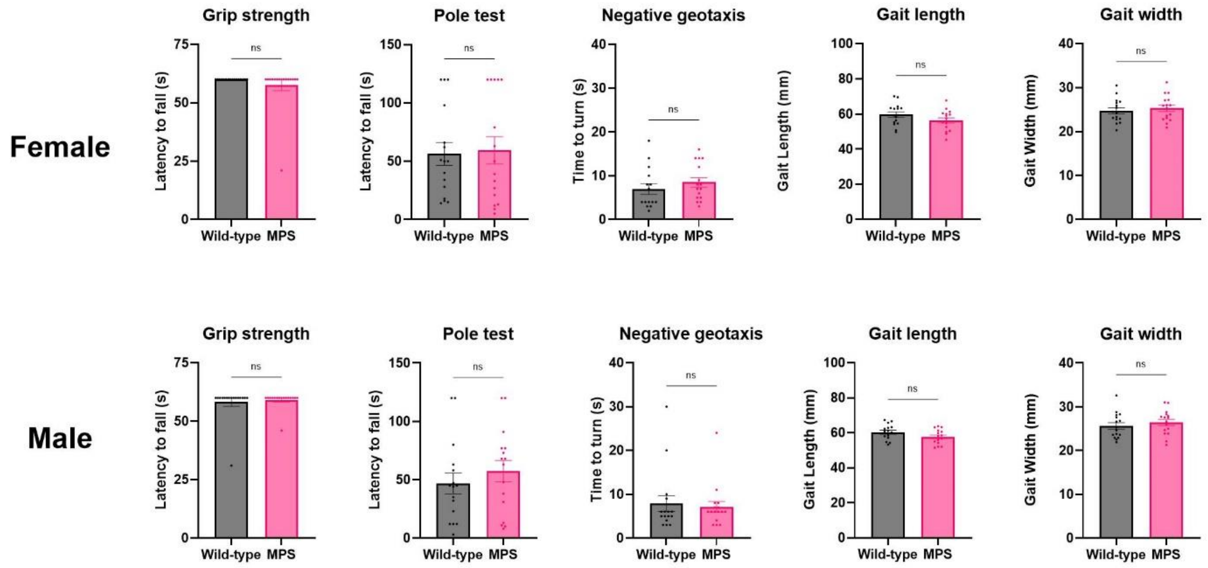


**Supplementary Figure 17: A gross motor phenotype is not apparent in MPS IIIA mice at 25 weeks of age.** Female and male wild-type and MPS IIIA were evaluated in a series of motor function tests on consecutive days. Neuromuscular grip strength was unchanged between genotypes. In the pole test, the time taken to turn 180 degrees and climb down a pole was measured. In the negative geotaxis test, the time taken to re-orient their body 180 degrees to face towards the ceiling was measured. The walking pattern of the hind paws was measured from at least two replicate tracks. The hind-limb gait length and width were measured from at least three strides per track. n=16 mice/group. ns, not significant.
